# Tandem repeats in methanogenic archaea and their viruses reveal a shared potential for rapid genomic diversification

**DOI:** 10.64898/2026.10.30.763056

**Authors:** Sebastian Haiß, Emilie J. Skoog, Josephine Herbert, Chang Ding, Katrin Weidenbach, Muhammad Aammar Tufail, Sylvia Schlaich, Bastian Molitor, Ruth A. Schmitz, Marie C. Schoelmerich

**Affiliations:** Environmental Microbiology, Department of Environmental Systems Science, ETH Zürich, 8092 Zürich, Switzerland; Department of Molecular Environmental Biotechnology, Helmholtz Centre for Environmental Research - UFZ, 04318 Leipzig, Germany; Institute for General Microbiology, Kiel University, 24118 Kiel, Germany; Climate and Environmental Physics and Oeschger Centre for Climate Change Research, University of Bern, 3012 Bern, Switzerland; Microbial Metabolic Biochemistry, Institute of Biochemistry, University of Leipzig, 04103 Leipzig, Germany

## Abstract

Methanogenic archaea drive global methane emissions, yet the molecular basis of their physiological flexibility is unclear. Archaeal adaptation is typically attributed to horizontal gene transfer, which is episodic and cannot explain rapid, reversible change. Here we show that direct tandem repeats (TRs) offer an intrinsic alternative. By screening 50,567 prokaryotic genomes, we find TR densities in *Methanosarcina* more than 30-fold above the prokaryotic average, paralleling Borg extrachromosomal elements. These TRs recur at conserved loci while sharing almost no sequence identity between strains, indicating independent origin, and evolve during routine cultivation. Methanogen TRs are transcribed, show ribosome occupancy, and lie near regulatory and cofactor-related genes, whilst being depleted around core energy-conserving and translational machinery. Genic TRs generate intrinsically disordered regions in proteins carrying nucleotide-binding, protein-protein interaction and cell-surface domains, and occur in structural and surface proteins of methanogen-infecting viruses. TRs are therefore an overlooked source of variability and adaptation potential in methanogens and their viruses.

## Introduction

Methanogenic archaea (methanogens) are the only organisms capable of biological methane production. They inhabit anoxic ecosystems ranging from marine and freshwater sediments to permafrost, wetlands, rice paddies, intestinal tracts, landfills and anaerobic digesters, and account for ∼70% of global methane emissions^1^. Among them, *Methanosarcina* is exceptional in metabolic and morphological flexibility and the only genus encompassing all three methanogenesis pathways: hydrogenotrophic, acetoclastic and methylotrophic^2^. *Methanosarcina mazei* Gö1, the strain we focus on in this work, can utilize H_2_ + CO_2_, acetate, methanol and methylamines for growth, and extensively remodels its transcriptome when switching between substrates^3^. It also alternates between multicellular aggregates (referred to as sarcinae) and single cells depending on osmolarity^4^, a transition mediated by a secreted disaggregatase that degrades the methanochondroitin envelope^5^. However, the triggers that determine individual cell fate during this switch remain unknown.

Several adaptive mechanisms in methanogens are controlled by transcription factors^6^ and by an extensive repertoire of small regulatory RNAs (sRNAs) and small proteins^7–9^, yet other adaptive phenomena remain unexplained. The lytic virus MetSV infects only *Methanosarcina* growing as single cells, with resistance in *M. mazei* arising through selection of the aggregate morphotype from a heterogeneous culture rather than through CRISPR-Cas spacer acquisition^10^, pointing towards a pre-existing, heritable variation within clonal populations. Heritable change also arises during routine passage, such as when *M. barkeri* Fusaro acquired stable fast growth on pyruvate after a prolonged lag^11^. Archaeal innovation is generally attributed to horizontal gene transfer (HGT)^12^. In fact, acetoclastic methanogenesis in *Methanosarcina* is thought to have been acquired from cellulolytic Clostridia^13^, and the original genome analysis assigned one third of the *M. mazei* Gö1 gene repertoire a bacterial ancestry^14^. But HGT is episodic, largely irreversible, and requires donor DNA, so it cannot explain rapid, reversible, strain-specific variation within clonal populations.

One mechanism that can generate rapid, reversible genetic variation is unit number change at perfect direct tandem repeats (TRs). TRs are mutational hotspots that expand and contract via replication slippage and recombination, driving rapid evolution and phenotypic diversification in eukaryotes and bacteria^15^. In eukaryotes, TRs modulate transcription, influence chromatin organization, serve as rapidly evolving cis-regulatory elements, facilitate *de novo* gene birth, and trigger repeat-expansion diseases when destabilized^15^. Within open reading frames (ORFs), they frequently encode intrinsically disordered regions (IDRs) that mediate interactions with DNA, RNA, proteins and lipids^16^. In bacteria and fungi, TR-driven phase variation enables reversible switching of gene expression to remodel cell-surface structures or evade host immunity^15,17^. In archaea, by contrast, the role of TRs in generating variation has received little attention, and because no pathogenic archaea are known^18^, it is unclear whether TR-mediated variation plays a comparable role.

Scattered reports indicate that TRs exist in methanogens^19^, but a systematic investigation has been lacking: a plasmid-borne *orc1* gene in *M. barkeri* is flanked by a putative origin of 38 TR units^20^; archaeal histones assemble on trinucleotide TRs^21^; amino acid repeats occur in cell-surface proteins with cryptic function^22^; and tracts in the *vho* hydrogenase operon and arginine biosynthesis pathway were proposed to be regulatory elements^23,24^. Borg extrachromosomal elements associated with “*Candidatus* Methanoperedens” (*Methanoperedens*) brought the question into focus: Borg genomes carry pervasive TR tracts that evolve rapidly and arose independently, whereas TRs are virtually absent from host genomes^25^. However, neither Borgs nor *Methanoperedens* have been obtained in pure culture, which has precluded experimental investigation.

We therefore searched for analogous TR occurrence and dynamics across prokaryotes. Combining comparative genomics, long-read sequencing, transcriptomics, ribosome profiling and proteomics, we show that *Methanosarcina*, close relatives of *Methanoperedens,* and three gut-associated methanogen genera harbor TRs paralleling those in Borgs. Focusing on *M. mazei* Gö1, we resolve TR dynamics within a single population and establish that TRs are transcribed and translated *in vivo*. Finally, we extend the analysis to methanogen-infecting viruses, suggesting that TRs may also drive adaptation in archaeal host-virus systems.

## Results

### Methanogen genomes harbor pervasive nucleotide tandem repeats

TRs are pervasive in Borg genomes, where they are proposed to drive diversification^25^. Given this, we asked how widespread TRs are beyond Borgs, surveying their occurrence across the prokaryotic tree of life (TRs defined as ≥ 3 identical units spanning ≥ 50 bp). Screening 50,567 genomes (49,948 bacterial, 619 archaeal; Supplementary Table 1) revealed exceptional densities in certain methanogen lineages: all 30 *Methanosarcina* genomes from a wide range of anoxic habitats including sewage digesters, marine and lake sediments, rice paddy soils, peat bogs, groundwater, and bovine rumen content (Supplementary Table 1) harbored 33–334 TR regions across genome sizes of 3.1–5.7 Mb at a mean of 4.32 ± 2.16 TRs per 100 kb, 35-fold above all other prokaryotic genomes (0.12 ± 0.24 TRs per 100 kb, n = 50.537; Wilcoxon rank-sum test, *p* = 2.3 × 10^-21^) and closely mirroring Borgs (5.10 ± 1.75 TRs per 100 kb, n = 17; Fig. 1).

**Fig. 1.**
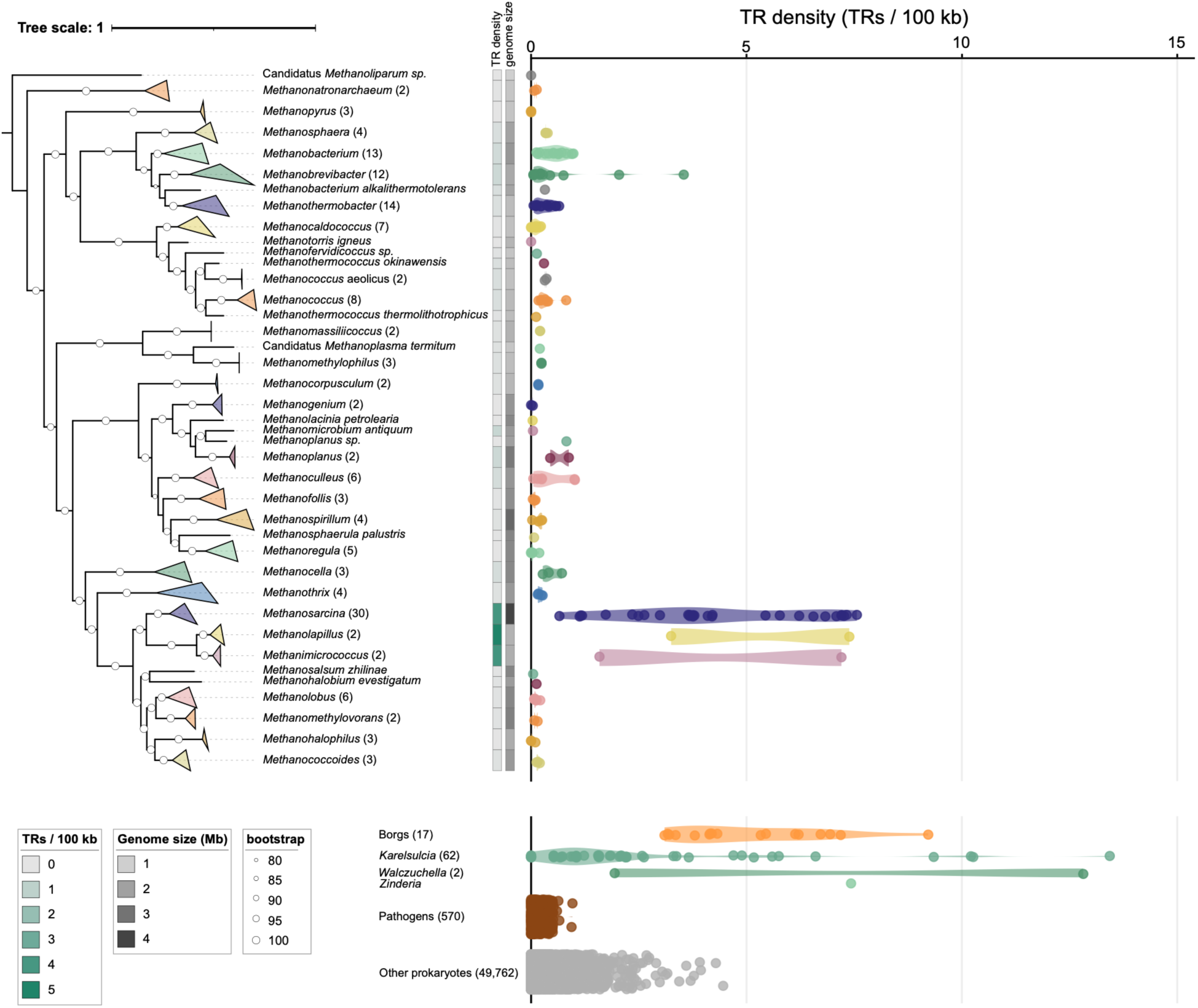
Comparative analysis of tandem repeat (TR) density across prokaryotic genomes and Borg extrachromosomal elements. TR density of methanogen genomes (n = 170), Borgs (n = 17), bacteriocyte-associated endosymbionts of sap-feeding insects (*Karelsulcia*, *Walczuchella* and *Zinderia*; n = 65), known TR-bearing bacterial pathogens (*Neisseria* and *Haemophilus*; n = 570), and all other prokaryotes (n = 49,762) from the RefSeq database. Number of genomes per genus and group is shown in parentheses.

Three further taxa displayed comparably high densities, all associated with animal digestive tracts: the arthropod gut-associated *Methanolapillus* (5.32 ± 2.93 TRs per 100kb, n = 2) and *Methanimicrococcus* (4.40 ± 3.97, n = 2), and the rumen-associated *Methanobrevibacter ruminantium* (3.54, n = 1) and *M. olleyae* (2.04, n = 1). Rumen and arthropod hindgut are both dense, host-structured anoxic environments dedicated to the degradation of recalcitrant plant polymers, in which methanogens depend on fermentation products supplied by a fluctuating microbial community and face intense viral predation. Since no plant polysaccharide-degrading enzymes were detected in methanogen genomes (Supplementary Table 2.1) and methanogens are not known to degrade plant polymers directly, the parallel between these lineages is ecological, resting on syntrophic dependence within a heterogeneous community. In contrast, the human gut-associated *M. smithii* harbored very few TRs (0.12 ± 0.09, n = 5), so elevated TR density is neither universal among methanogens nor among host-associated ones. Phylogenetic lineage (*p* = 3.17 × 10^-15^) and genome size (*p* = 1.96 × 10^-5^) were the primary drivers of TR density, whereas ecosystem classification, predicted growth conditions, and encoded methanogenesis pathways showed no significant effect.

Among non-methanogens, elevated densities were observed in sap-feeding insect endosymbionts *Karelsulcia*, *Walczuchella* and *Zinderia* (2.51 ± 3.05, n = 65; Fig. 1), where extreme genome reduction inflates any per-100-kb metric^26^. Notably, *Neisseria* and *Haemophilus*, the best-characterized TR-bearing pathogens^17^, showed low TR densities under our stringent perfect-repeat criteria (0.17 ± 0.15, n = 570). This positions *Methanosarcina* as an exception among free-living prokaryotes.

### *Methanosarcina* TRs maintain compositional signatures despite sequence divergence

Across 87 dereplicated (95% ANI) methanogen genomes, TR regions were short (median 75 bp, four units; up to 1,428 bp or 22 units; Supplementary Table 2.1 and 2.2; Supplementary Fig. 1), accounting for a median of 0.07% of genome length. Averaged across genomes, 55.1 ± 34.6% of TRs were intergenic, 39.1 ± 35.2% were genic and 5.8 ± 15.6% overlapped ORF boundaries (n = 83 genomes). Compared to a mean coding density of 82.2 ± 7.6%, intergenic regions were disproportionately TR-enriched, and genic TRs preserved the reading frame in 90.8% of cases. TRs were enriched in adenine and thymine (+6.4% and +14.0%; p_adj_ < 2.2 × 10^-16^), and depleted of guanine and cytosine (−11.0% and −25.1%; p_adj_ < 2.2 × 10^-16^; Supplementary Fig. 2), the same signature as Borg TRs^25^.

Within *Methanosarcina* these trends were stronger (77.0% intergenic; A: +16.1%, T: +16.6%, G: −22.0% and C: −24.9%; p*_adj_* < 2.2 × 10^-16^; Supplementary Table 3). Despite the shared signature, unit sequences were highly diverse, with fewer than 2% of TRs shared within a genus (Fig. 2A), fewer than 25% within *M. mazei* (Supplementary Fig. 3; Supplementary Table 4.1), and TRs with identical units across strains often differed in unit number (Supplementary Table 4.2). Individual regions were either conserved in sequence and unit number, conserved in sequence but variable in unit number, or collapsed to a single non-repetitive unit (Fig. 2B). In the S-layer and Cdc6 cases the TR lies beside an accessory paralog rather than the dominant copy, suggesting repeat variability is tolerated where it is unlikely to be lethal.

**Fig. 2.**
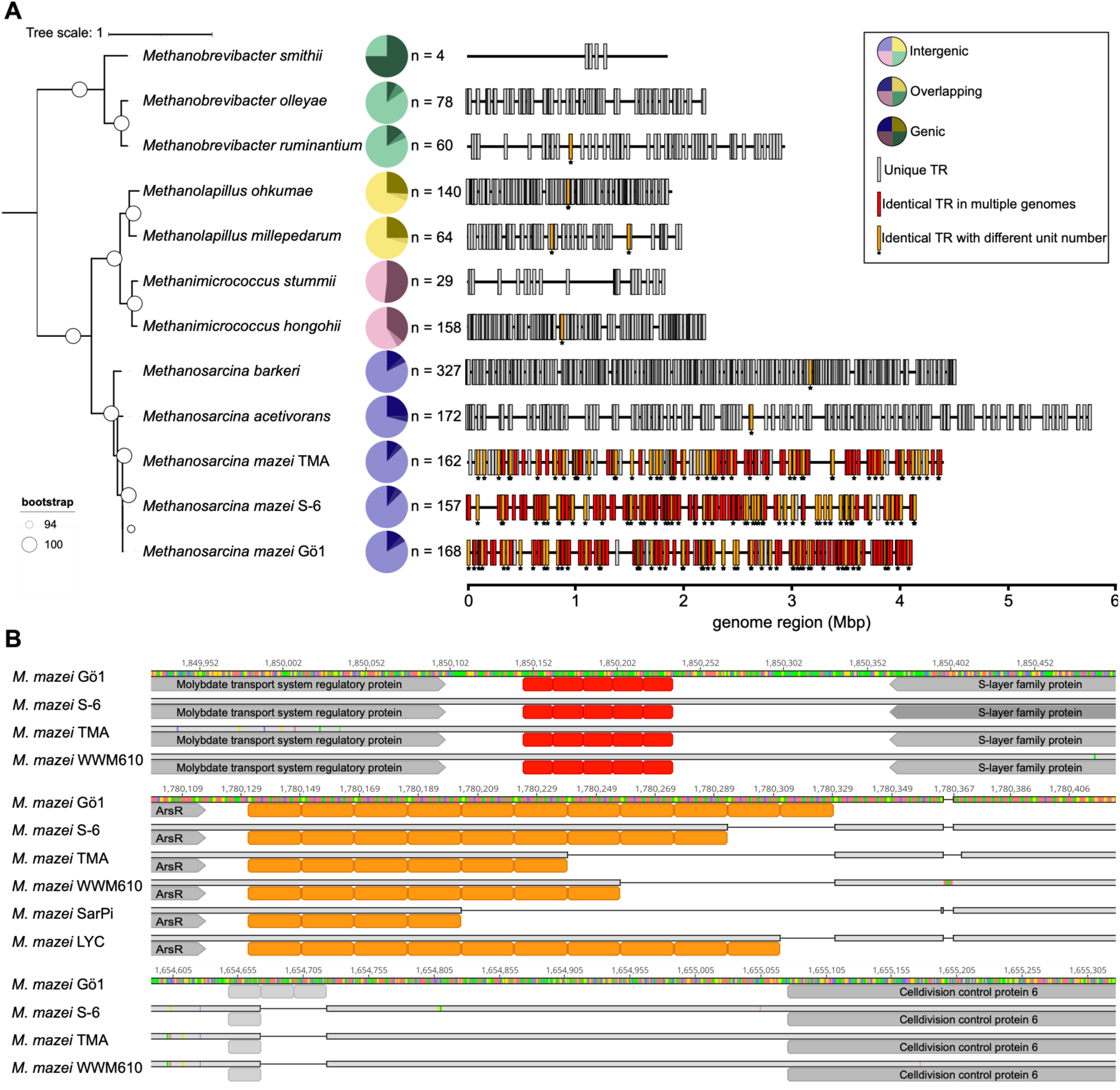
Diversity and genomic distribution of TRs in methanogen genomes. **A** Phylogenetic tree of selected methanogens showing absolute TR counts and the proportion of intergenic, genic, and ORF-overlapping TRs (pie chart). TRs are shown on the genome color-coded to distinguish unique TRs (grey) from TRs shared across multiple of the represented genomes (red) and shared TRs with unit number variation (orange, asterisks). **B** Examples of TR dynamics across *M. mazei* genomes, showing a TR unit number-conserved TR (red) adjacent to the S-layer gene, a TR with conserved sequence but variable TR unit number (orange) adjacent to a regulator gene, and a strain-specific TR absent from other genomes (grey) adjacent to a Cdc6 gene. Nucleotide sequences of *M. mazei* Gö1 are displayed with red (A), green (T), yellow (G), and blue (C).

### TRs evolve rapidly within a *Methanosarcina mazei* population

To assess variability within a single strain, we performed long-read whole-genome sequencing (WGS) of *M. mazei* Gö1 cultures from DSMZ (DSM3647), Kiel University (Ref_WT), and ETH Zürich (Ref_ETH; see Methods). All three were nearly identical to one another and to the reference genome (NC_003901) by ANI (Supplementary Fig. 4), yet their TR regions had already diverged. Of the 168 reference TRs, two were absent from Ref_ETH strain due to a 23 kb genomic deletion. Ref_WT and Ref_ETH had each lost one repeat unit at TR_115 and TR_125 and carried single nucleotide variants (SNVs) within TR_82 and TR_131. TR_125 overlaps the 5′-end of a helix-turn-helix transcriptional regulator gene (MM_RS14810); TR_115 lies downstream of a ferritin-like hypothetical protein gene (MM_RS14225). All cultures had gained repeats: one region in DSMZ and four in each of Ref_WT and Ref_ETH, of which two were caused by reference SNVs and two reflected true unit gains (Supplementary Tables 5.1 and 5.2). Both newly expanded TRs lie close to existing intergenic TR regions (TR_13 and TR_146), identifying these loci as mutational hotspots that continued to evolve during routine laboratory maintenance in liquid media.

Read-level heterogeneity in unit number differed strongly between regions (Fig. 3A). The most variable, TR_24, deviated in 4.6% of reads (up to 9.4%), gaining up to three and losing up to four units (Supplementary Fig. 5), and lies just upstream of the mapped 5′-UTR of the glycogen synthase gene (MM_RS03065; Fig. 3B). Since *M. acetivorans* uses glycogen as a carbon reserve^27^, 5′-UTR length variation could influence glycogen synthesis. TR_123 (3.0% of reads, up to 6.2%) lies between an APC-family transporter (MM_RS14765) and a SET-domain lysine methyltransferase (MM_RS14760; Fig. 3C), a class that modulates chromatin accessibility in eukaryotes^28^. TR_83 (1.5%) lies 1,643 bp upstream of the CODH/ACS complex, which cleaves acetyl-CoA during acetoclastic methanogenesis, too distant for direct regulation but potentially affecting long-range DNA structure (Fig. 3D); TR_8 (1.1%) spans almost an entire ORF overlapping a tRNA pseudouridine synthase gene, an RNA-binding enzyme with possible chaperone function^29^, so variation should affect both transcripts (Fig. 3E).

**Fig. 3.**
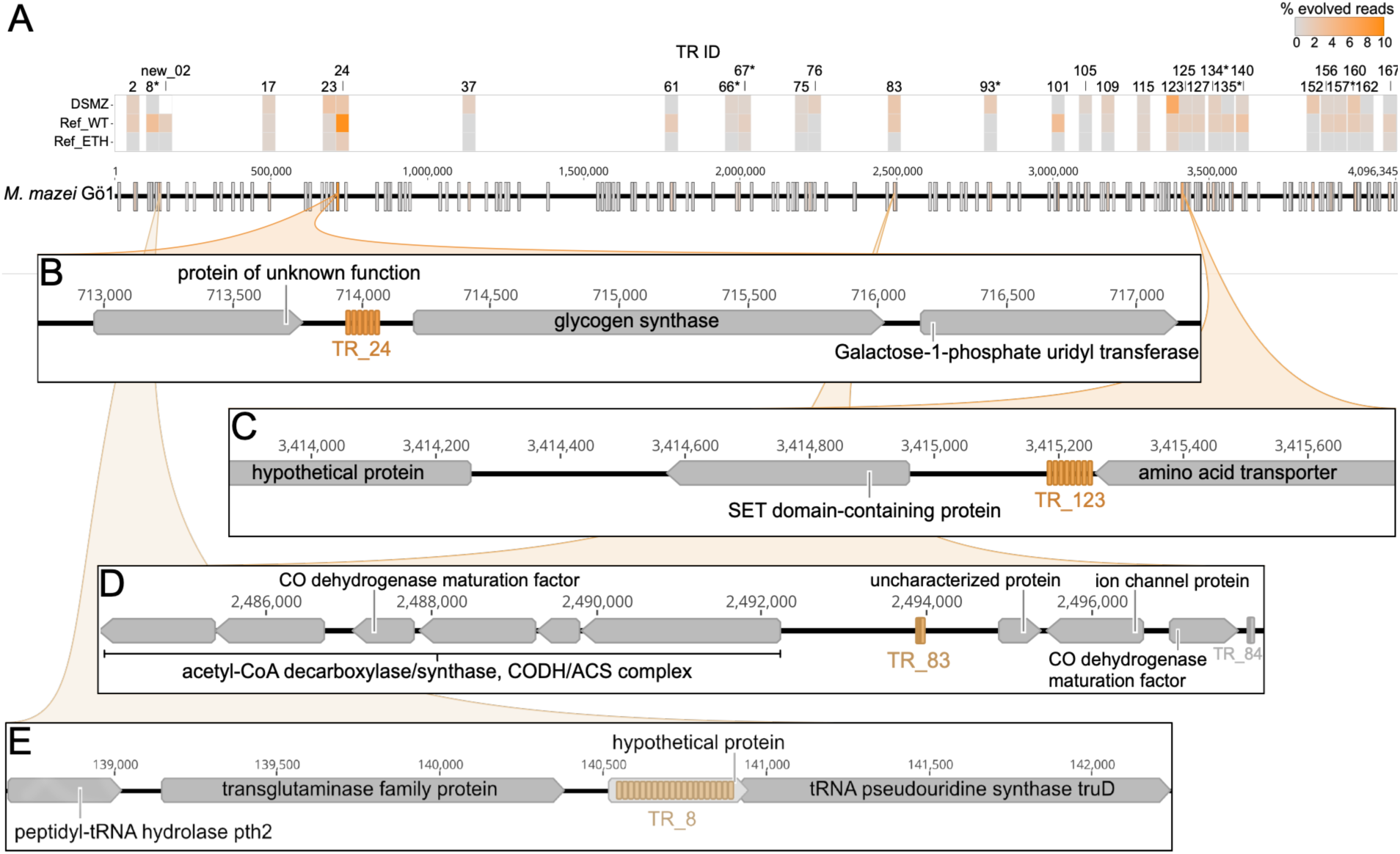
Within-population heterogeneity of TR unit numbers in *M. mazei* Gö1. **(A)** Heatmap showing the proportion of reads deviating from the population mean unit number for the 20 most variable TRs across three independently propagated cultures (DSMZ, Ref_WT, Ref_ETH). Asterisks mark genic TRs. **(B-E)** Detailed view of the genomic neighborhood for the four most dynamic TR loci in detail: **(B)** TR_24, upstream of a glycogen synthase gene; **(C)** TR_123, between an APC-family transporter and a SET-domain protein; **(D)** TR_83, upstream of the CODH/ACS complex; **(E)** TR_8, within an ORF adjacent to a tRNA pseudouridine synthase.

Unit number changes occurred in genic, ORF-overlapping and intergenic TRs, with no significant difference in magnitude between the three classes (Kruskal-Wallis test, *p* = 0.059). For comparison, the same read sets contained 15–190 SNVs per genome at mean allele frequencies of 25.0–41.0% (Supplementary Table 6). Thus, whereas point mutations were distributed across the genome, TR unit-number variation was concentrated at a few loci at which up to ∼9% of reads deviated from the consensus.

### TRs lie near genes with regulatory and cofactor-related functions

We next examined which neighboring genes might be affected. Of 85,556 genes across 28 *Methanosarcina* genomes, 6,209 genes (7.26%) lay adjacent to one of the 3,270 TRs. Several KEGG Orthology level C categories were significantly overrepresented among TR-neighboring genes: peptidases and inhibitors, two-component systems, metabolic pathways (pyruvate, pantothenate/CoA, selenocompound/ascorbate/aldarate), and genetic information processing (tRNA biogenesis, chromosome-associated proteins; Extended Data Fig. 1A, Supplementary Table 7). Conversely, prokaryotic defense systems, transcription machinery, ribosomes, and core energy metabolism, including methane metabolism, were significantly underrepresented. *Methanosarcina* therefore maintains TRs near specific adaptive and regulatory pathways while shielding core energy-generating and translational machinery from TR-driven variability.

This exclusion is not absolute. For a subset of genes, more than 40% of copies lay adjacent to a TR (Extended Data Table 1), including glycerol-1-phosphate dehydrogenase (K00096), which produces the *sn*-glycerol-1-phosphate backbone that defines archaeal membrane lipids^30^; tyrosine/aspartate decarboxylase (K18933), of methanofuran biosynthesis; and the replication initiator Cdc6/Orc1 (K10725). Two further hits, methylthioribose-1-phosphate isomerase (K08963, *mtnA*) and methionine synthase (K00549, *metE*), constrain the supply of methionine and thus *S-*adenosylmethionine (SAM), potentially affecting SAM-dependent methylation and radical SAM chemistry.

Even where the same gene was recurrently TR-associated, the TRs differed markedly in sequence, length, and position (Supplementary Fig. 6). A striking example is *ftr*, encoding the formylmethanofuran-tetrahydromethanopterin *N*-formyltransferase of the second methanogenesis step^2^ (Extended Data Fig. 1B and C). Although methanogenesis genes are globally TR-depleted (*p_adj_* = 0.024), TRs flanked *ftr* in 38% of the genomes in which it was identified (n = 29). Their retention despite genome-wide depletion suggests a specific benefit, and the absence of conservation in position, sequence, and length suggests that what matters is a variable-length tract rather than identity.

### Many TRs are transcribed and respective RNAs show ribosome occupancy

Published RNA-seq and Ribo-seq data from *M. mazei* Gö1 under nitrogen sufficiency or stress^7^ showed RNA-seq coverage at 59 of the 168 TRs (13 moderate, 3 high) and Ribo-seq coverage at 63 (11 moderate, 9 high; Supplementary Table 8 and Supplementary Fig. 7). Because raw read counts cannot separate occupancy from transcript abundance, these data provide evidence of occupancy across TR-containing transcripts rather than translation of the tracts themselves. Seven TRs with moderate to high signal lay within ORFs encoding the chemotaxis histidine kinase CheA (MM_RS06875; Supplementary Fig. 7D-F), two transcriptional regulators (MM_RS08605, MM_RS08635), a cell adhesion-related protein (MM_RS15745), a 5-oxoprolinase (MM_RS15110), an extracellular lactonase domain containing protein (MM_RS15170), and a hypothetical protein (MM_RS10295) that is 87% TR sequence and predicted fully disordered. This set is dominated by signal transduction (CheA), transcriptional regulation (two regulators) and surface adhesion – the same categories enriched genome-wide among TR-adjacent genes, and precisely those for which reversible, tunable variation would be advantageous.

Two expressed TRs overlapped the small RNAs, sRNA_75_ and sRNA_129_^31^. Long-read transcriptomics captured full-length transcripts with sharply defined boundaries (Extended Data Fig. 2) at levels above the constitutively high *mcr* genes used as an internal reference (Supplementary Fig. 7G). sRNA_129_ additionally showed a Ribo-seq coverage, consistent with a potential dual function as regulatory RNA and small protein, although coverage dropped across the TR region itself. Both sRNAs consist predominantly of TR sequence, so changes in unit number will likely directly alter transcript length, secondary structure, stability, and target binding. IntaRNA predictions identified candidate mRNA targets dominated by genome-rearrangement functions, including transposases and a Cas1-solo likely to be a casposon transposase (Supplementary Table 9), though these rest on hybridization energy alone and require experimental validation.

### Genic TRs encode disordered extensions concentrated in cell-surface proteins

Genic TRs form amino acid tandem repeats (aaTRs) that are rapidly evolvable protein regions. *Methanosarcina* genomes encode 34.3 ± 16.9 TR-containing proteins each (n = 960), typically one repeat region per protein (1.15, n = 1,102) of 29.9 ± 17.9 residues built from 5.0 ± 2.9 units of 6.4 ± 4.0 residues. 110 proteins harbored multiple aaTRs and extracellular TR-proteins were enriched for multi-TR architectures (*p* = 0.003). Mirroring the nucleotide bias, aaTRs were enriched in serine, asparagine, threonine and glutamic acid and depleted of glycine, arginine, histidine, and all hydrophobic or aromatic amino acids (Fig. 4A), yet diverged between homologs (Extended Data Table 2) and sub-cellular localizations (Fig. 4B and 4C). The exosome component RRP41^32^ exemplifies this (Fig. 4D): its N-terminal domains are conserved while its C-terminal aaTR is variable, disordered and glutamate-dominated, carrying exclusively synonymous SNVs (Supplementary Fig. 8).

**Fig. 4.**
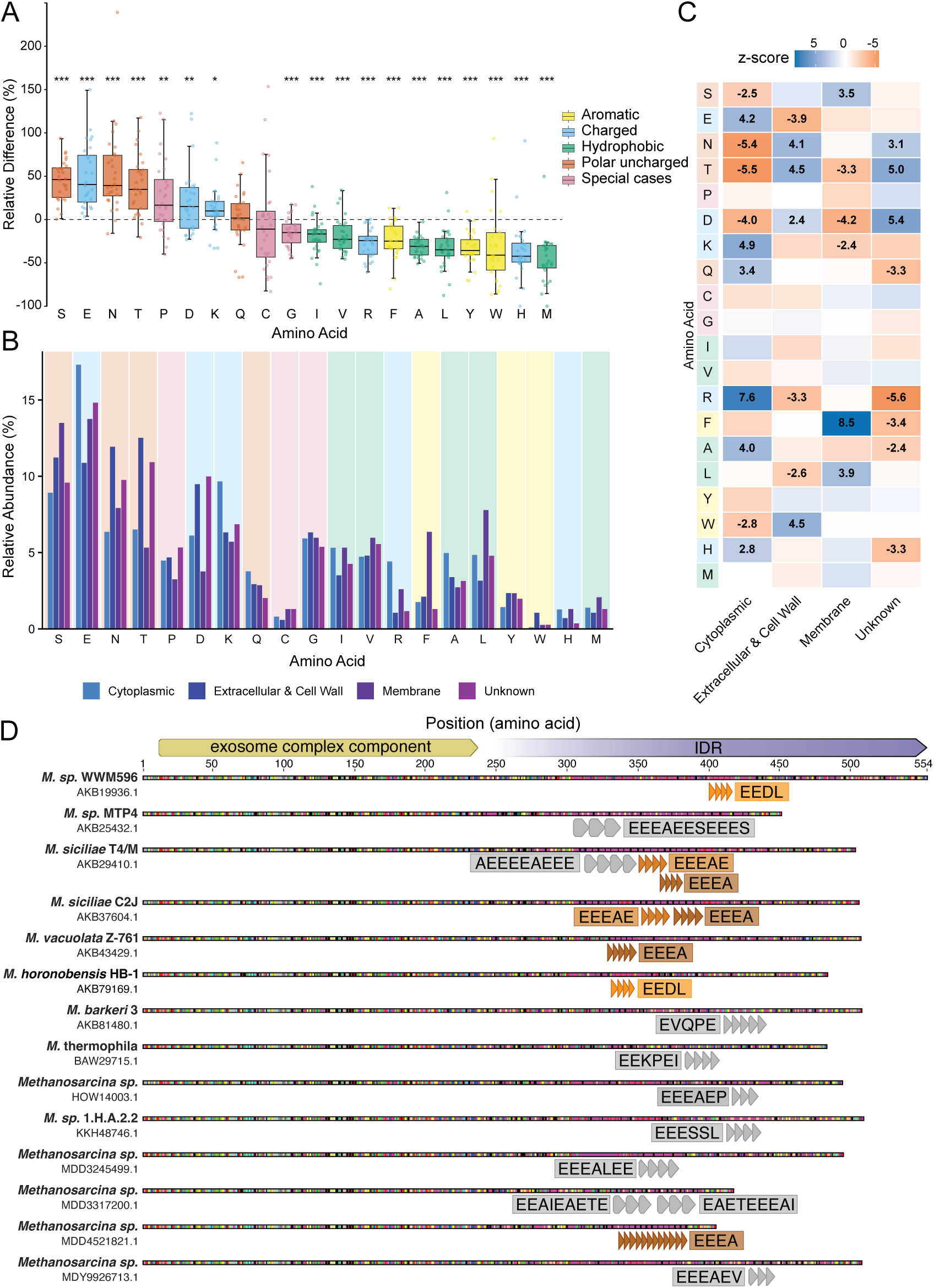
Properties of amino acid tandem repeats (aaTRs) encoded by *Methanosarcina* genomes. **(A)** Relative difference in amino acid abundance of aaTRs compared to the background proteomes. Colors indicate amino acid physicochemical properties. **(B)** Relative amino acid abundance in aaTRs, stratified by predicted subcellular localization. **(C)** Corresponding amino acid enrichment and depletion patterns expressed as standardized residuals (*Z*-scores), where positive values denote enrichment and negative values denote depletion. **(D)** Alignment of exosome complex component RRP41 sequences and their aaTRs (triangles) of *Methanosarcina* spp. Corresponding aaTR sequences are displayed adjacent to the TRs, with identical aaTR sequences highlighted in matching shades of orange. The propensity for predicted intrinsic disorder regions (IDR) increases towards the C-terminus.

Of 95,121 *Methanosarcina* proteins, 67,661 (71.1%) could be functionally predicted, as could 504 of the 960 aaTR-proteins (52.5%). TR-proteins were enriched in motility, mRNA biogenesis, DNA replication, chromosome-associated processes, and peptidase/inhibitor activity (*p* < 0.007; Supplementary Table 10), and absent from nucleotide metabolism, glycerophospholipid metabolism, and signaling. Repeats also occur in DP1, the proofreading subunit of the archaea-specific D-family polymerase (PolD; K02323)^33^, and in CheA, which interfaces with the archaellum^34^. Protein clustering placed 90.7% of all 95,121 proteins into 7,228 clusters, and the aaTR-richest were dominated by surface proteins (Extended Data Table 2): the top two are extracellular or cell-wall associated, carrying signal peptides, IDRs and PKD (IPR035986) or cell-adhesion domains (CARD; IPR011635).

Functional domains in TR-proteins were enriched for nucleotide binding and protein-protein interaction (*p_adj_* < 5.5 × 10^-05^), with aaTRs almost always spatially separated from them (Supplementary Table 11), suggesting repeats extend proteins without disrupting their core function. TR-proteins were predicted 2.9 times more often in the cell wall and 6.8 times more often extracellularly than the overall proteome (*p* = 9.99 × 10^-05^). Among predicted cell-surface proteins, the most strongly enriched domains included the disaggregatase-related domain (107/201 proteins, *p_adj_* = 1.83 × 10^-115^) and the disaggregatase-related repeat domain (84/114 proteins, *p_adj_* = 1.32 × 10^-108^; Supplementary Table 12). Although perfect aaTR co-occurred with these in only six and one proteins, respectively, repeat-level variation was common at a larger scale, as of 257 disaggregatase-related proteins, 122 contained a repeat domain one to three times (Supplementary Table 13), and 57 carried degenerate repeats whose unit number could change through recombination^35^.

Most strikingly, TR-proteins were strongly enriched in IDRs (*p* = 4.02 × 10^-290^), with the TR nested directly within the IDR in 85.0% of cases, mirroring Borg-encoded aaTRs^25^. Although TR-IDR association is documented across all domains of life^16^, our data suggest that in *Methanosarcina* repeats frequently generate IDRs rather than coincide with pre-existing disorder. Outside the genus *Methanosarcina*, *M. ruminantium* (NC_013790.1) showed the same surface bias (17 perfect and 26 degenerated aaTRs, 20.9% surface-localized, Supplementary Table 14), including an adhesin-like protein (WP_048812472.1) that mediates binding to H_2_-supplying protozoa and is a candidate vaccine target^36^.

Analysis of published whole-proteome datasets detected 45.5%, 15.9%, and 50.0% of TR-proteins in *M. mazei* Gö1, *M. acetivorans* C2A, and *M. olleyae*, respectively. Only one peptide lying within an aaTR passed the 1% false discovery rate (FDR) threshold (‘ILELETLLVDEQR’ of *M. acetivorans* AAM05742.1; peptide-level q-value < 0.001, four peptide-spectrum matches). Over half of the aaTRs in these TR-proteins lack lysine and arginine and therefore lie within tryptic peptides longer than the 30 residues searched; in most of the others, closely spaced cleavage sites yield searchable peptides only through missed cleavages, if at all. Nevertheless, five of the seven *M. mazei* ORFs containing a TR with moderate to high Ribo-seq signal were detected at the protein level (Supplementary Table 15A), so TR-proteins are produced *in vivo*, although peptide-level evidence for the repeat tracts is so far limited to the *M. acetivorans* peptide noted above.

### TRs occur in viruses that infect methanogens

Borgs are exceptionally rich in TRs while their *Methanoperedens* hosts carry virtually none^25^, so we asked whether methanogen extrachromosomal elements also carry TRs. Among nine published methanogen plasmids, TRs occurred in only two, both from *M. barkeri* (Supplementary Fig. 9 and Supplementary Table 16), in each case adjacent to the replication origin and Cdc6 initiator. This is consistent with previous reports of repeats forming bubble-out structures that facilitate replication initiation^20^ and mirrors the observed TR occurrence next to Cdc6 in *Methanosarcina* chromosomes. Among isolated methanogen viruses, only ΨM2, which infects *Methanothermobacter marburgensis* and establishes either a lytic or a non-lytic lifecycle,^37^ harbored a TR within the gene encoding a tail tape measure protein (TMP; Supplementary Table 17).

Mapped reads from a recent ΨM2 study^37^ revealed that the TMP repeat exists in multiple length variants: 44.6% of reads deviated from the consensus of 13 units, spanning a continuous range from complete loss (2 units) to 28 units, skewed towards contraction (Fig. 5A). This has a direct consequence for domain calling. At the 10 copies present in the reference (NC_001902.1), the PKIEF-based aaTR matches no annotatable domain, whereas above 12 copies the same region is annotated as a mucin-2 domain (PTHR33088), demonstrating that TR expansion can directly give rise to a novel protein domain (Supplementary Fig. 10). Since TMP length determines tail length and the TMP sequence has been implicated in host-range determination in tailed phages^38,39^, this variation may enable the virus population to broaden or shift its host range.

**Fig. 5.**
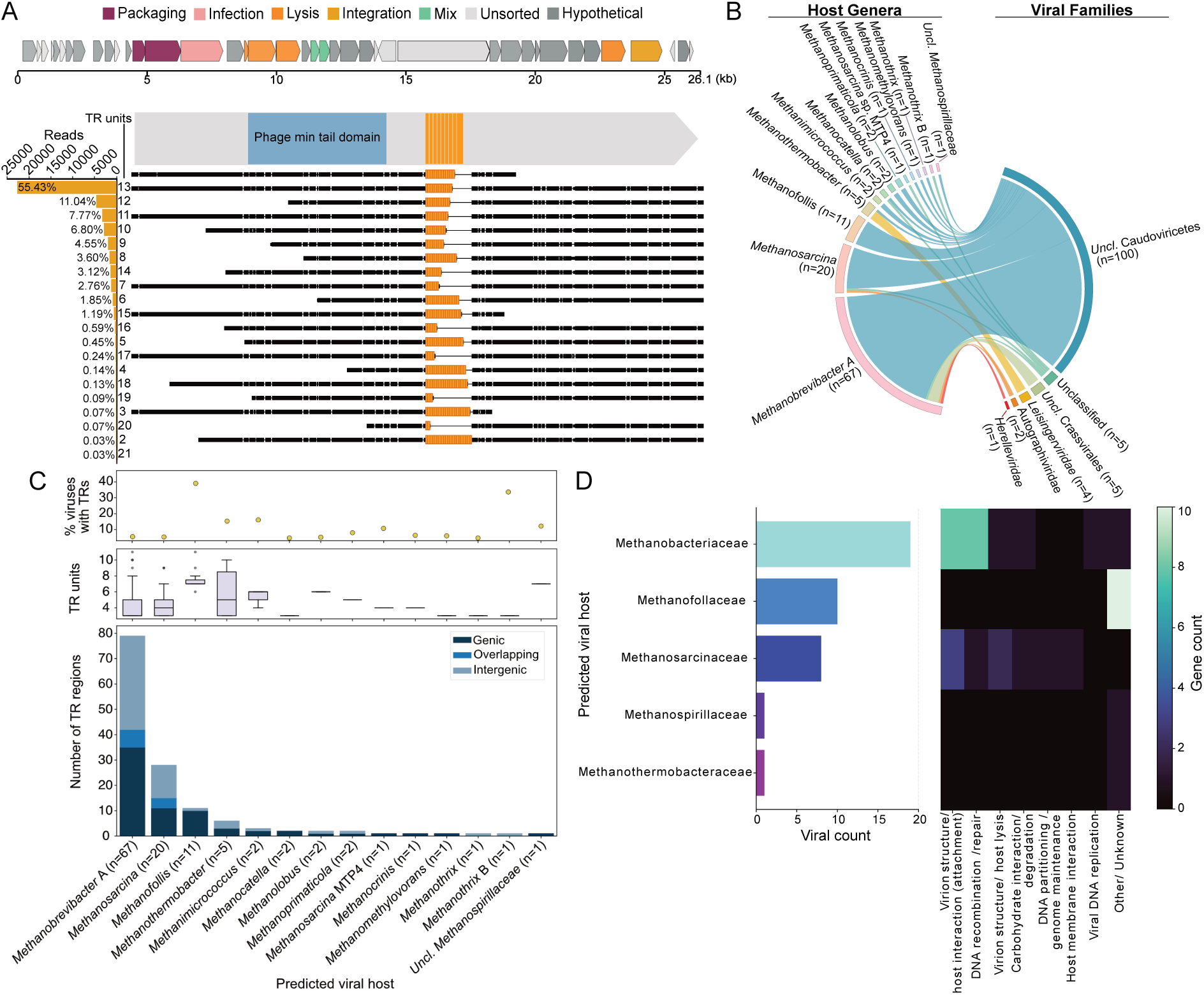
Genome-wide analysis and functional characterization of tandem repeats in methanogen viruses. **(A)** The genome of the *M. marburgensis*-infecting ΨM2 virus encodes a TR-containing tape measure protein that harbors a TR with pronounced variability in TR unit number. **(B)** Sankey plot showing viral families with TRs and their predicted methanogen hosts. **(C)** Viral families clustered according to their predicted methanogen hosts. The upper panel displays the percentage of viral genomes containing at least one TR, the middle panel shows the distribution of TR units per TR region, and the lower panel highlights the proportions of genic, ORF-overlapping, and intergenic TRs. **(D)** Heatmap of gene counts showing viral TR-proteins clustered by predicted host family and functional annotation.

Across a global dataset of 407,171 metagenome-derived viral genomes (vMAGs), 5,046 vMAGs passed quality filtering and 2.3% (n = 117) contained at least one TR region. These spanned predominantly unclassified Caudoviricetes (85.5%) and targeted a diverse set of methanogen hosts (Fig. 5B). Normalized per host genus, TR prevalence was highest for *Methanofollis* (39.3%; 11/28), *Methanothrix B* (33.3%; 1/3), and *Methanimicrococcus* (22.2%; 2/9), and lower for more highly represented genera including *Methanobrevibacter A* (2.2%; 67/3,032), *Methanosarcina* (2.9%; 20/682), and *Methanocatella* (0.6%; 2/333; Fig. 5C). Of 139 vMAG TRs, 59 were intergenic and 80 were ORF-associated. Unlike TRs in *Methanosarcina*, they were more often genic (49.6%, Fig. 5C), mirroring Borgs (44%)^25^. The TRs in vMAGs were concentrated in genes encoding structural and surface-exposed proteins, predominantly receptor-binding tip proteins and the DNA repair/recombination protein RadA (Fig. 5D and Supplementary Table 18). TRs in methanogen viruses evolve rapidly at the population level (as exemplified by ΨM2), are preferentially located in surface proteins mediating host-virus interactions and generate diversity consistent with ongoing virus-host co-evolution.

## Discussion

Methanogenic archaea occupy anoxic environments that differ widely in substrate, salinity, temperature and viral pressure, yet how a conserved genomic core supports that range has been unclear^40^. Archaeal adaptation is attributed largely to horizontal gene transfer^12^, which is infrequent and irreversible, or to point mutations, which lack the reversion frequency required for rapid switching. We find that perfect direct tandem repeats provide an intrinsic alternative that requires no donor DNA. Their density in *Methanosarcina* approaches that of Borg elements and is without parallel among free-living prokaryotes. Expanding and contracting reversibly, they generate genetic heterogeneity within populations and may operate at three levels: reshaping DNA architecture, affecting sRNA-based regulation and generating disordered protein extensions.

The most direct evidence that these tracts are active rather than relic comes from within a single population, where up to 9% of reads at the most variable loci deviated from the consensus unit number, and from three separately maintained *M. mazei* Gö1 cultures that differ in repeat content despite ANI above 99.95%, consistent with observations between closely related Borg genomes^25^. Comparative genomics across *Methanosarcina* is consistent with this, as unit sequences are essentially unique to each locus, yet their genomic contexts recur across strains. At the *ftr* locus, a TR flanks this central methanogenesis gene in a third of genomes, yet differs in unit sequence, length and position in every instance. Rapid turnover of an ancestral tract could erase sequence similarity, but not produce this positional scatter, so we favor independent origination at these loci on timescales short enough to be observed during laboratory passage.

The view that archaea adapt mainly by gene acquisition^12^ is contested by diversity-generating retroelements in DPANN archaea and archaeal viruses^41^, yet TRs of this density and reach have been largely overlooked as drivers of adaptation. They do not compensate for a scarcity of mobile DNA, since *M. mazei* Gö1 encodes 102 transposase genes^14^. The mechanisms differ in kind rather than in frequency, as transposition is largely irreversible, whereas repeat expansion and contraction alter tract length reversibly, preserve the reading frame, and require no donor sequence.

At the DNA level, TRs supply both the mutational engine and a substrate for architecture: AT-rich tracts slip readily during replication^42^, archaeal histones assemble into nucleosome-like structures on short repeats^21^ and repeats flank the replication origin of *M. barkeri* plasmids^20^. At the RNA level they enter a regulatory layer on which this organism already relies, comprising 208 predicted sRNAs^31^, several functionally characterized^7,8^. The two most strongly expressed TRs constitute the bulk of sRNA_75_ and sRNA_129_, so TR unit gain or loss would be expected to alter transcript length, folding, stability and target affinity. At the protein level, repeats generate disordered extensions separate from functional domains, leaving catalytic and binding cores intact, as for Borg aaTRs^25^. Since repeat number within a disordered region can govern phase separation and dynamic intracellular compartmentalization in other systems^43,44^, and although condensates have not yet been described in archaea, we speculate that aaTR unit number may also tune the threshold at which they form in methanogens.

TR-proteins are enriched at the cell surface while TRs are depleted across ribosomes, transcriptional machinery, defense systems and central energy metabolism, mirroring classic bacterial contingency loci^45^. We propose that this is a candidate genomic basis for phenotypic switching in *Methanosarcina*, specifically the reversible transition between methanochondroitin-encased aggregates and single cells, which is tied to a disaggregatase carrying variable numbers of repeat domains^4,5^. Such variation could explain MetSV resistance arising through selection of an aggregated morphotype^10^, as repeat length in adhesins governs this phenotype in fungi^46^. Perfect TRs are rare within disaggregatase-related *Methanosarcina* proteins, and their repetitive architecture is instead defined by domain-level duplications, which change through recombination rather than replication slippage, extending the principle to a related mechanism. In viruses, TRs concentrate in receptor-binding tip proteins and in RadA, and in ΨM2 the tape measure protein ranged from 2 to 28 units within a single population, which may allow sampling of several host-range states at once^38,47^. Finally, the rumen methanogen *M. ruminantium* is a target for anti-methanogen vaccines based on surface antigens^48^, and the adhesin-like vaccine candidate WP_048812472.1 carries repeats, so candidate targets should be screened for repeat content and repeat-free epitopes prioritized^49^.

We applied conservative detection criteria, restricting the analysis to perfect TRs of at least 50 nt and three units, so the densities reported here are lower bounds. Permitting a single mismatch recovered 43 additional tracts in *M. mazei* Gö1 (168 increased to 211 TRs), and the length criterion excludes the short homopolymeric and tetranucleotide repeats that mediate classic phase variation, which accounts for the low counts in *Neisseria* and *Haemophilus*. It remains unresolved whether read-level heterogeneity reflects distinct subpopulations or variation among genome copies within polyploid cells^50^, and all regulatory targets and functional assignments remain predictions. Linking a shift in unit number causally to a phenotype will require chromosomal alteration of repeat tracts, feasible in genetically tractable *Methanosarcina*, together with characterization of TR-containing sRNAs and purified protein length variants.

Tandem repeats are therefore a pervasive and rapidly evolving feature of specific methanogen clades and their viruses. They are concentrated at the interfaces where these organisms meet their environment and depleted from the core machinery that sustains them, suggesting a previously overlooked source of genetic variability in microorganisms central to the global methane cycle.

## Methods

### Organism and cultivation

*M. mazei* strain Gö1 (Ref_WT) cultivated at Kiel University was further cultivated in house at ETH Zürich (Ref_ETH) and *M. mazei* strain Gö1 substrain DSMZ (DSM3647) was obtained from the Leibniz Institute DSMZ-German Collection of Microorganisms and Cell Cultures GmbH. The Kiel and the ETH cultures derive from the same parental stock and have been transferred (subcultured) independently since February 2024, up to 100 times; all subsequent experiments were started from cryo-stock cultures. Cultures were grown at 37 °C without rotation in 12-ml Hungate tubes or 120-ml serum flasks under an N_2_+CO_2_ (80:20 [v/v]) gas atmosphere in modified DSM medium 120c prepared using anaerobic techniques^51^ (full composition in Supplementary Note 1), buffered with 10.1 mM NaHCO_3_ (pH 6.9), supplemented with 0.37 mM L-cysteine and 0.38 mM Na_2_S, and supplemented with 75 mM methanol and 20 mM trimethylamine as energy and carbon sources. Growth was determined by measuring the optical density at 600 nm (OD_600_).

### Genome retrieval, dereplication and phylogeny

Genomes were downloaded from the National Center for Biotechnology Information (NCBI) RefSeq database on 12^th^ of November 2024, selecting bacterial and archaeal entries designated as complete genomes or complete chromosomes (50,567; Supplementary Table 1) for the initial analysis of tandem repeat density among prokaryotes. For methanogen genomes (identified based on taxonomy) isolation sources were extracted from NCBI and BacDive^52^ and categorized into ten ecosystem groups based on sample representation (n > 6 versus ‘others’; Supplementary Table 1), predicted optimal growth conditions were obtained using GenomeSPOT v1.0.1^53^, and encoded methanogenesis pathways were identified using DRAM v1.5.0^54^ with default parameters.

For downstream analyses, methanogen genomes were dereplicated using dRep v3.5.0^55^ at a 95% ANI threshold, yielding 87 representative genomes with 83 proteomes (Supplementary Table 2), which were screened for encoded carbohydrate-active enzymes using DRAM v1.5.0^54^ with default parameters. Genomes annotated as belonging to the genus *Methanosarcina* were downloaded separately on 9^th^ of September 2024, quality filtered for > 90% completeness and < 5% contamination using CheckM v1.2.2^56^ (135 high quality genomes), and further dereplicated at a 99% ANI threshold (37 genomes with 28 proteomes; Supplementary Table 3). Pairwise average nucleotide identity (ANI) among the 135 high-quality genomes was calculated using fastANI v1.3.4^57^ (Supplementary Table 4.1). Eleven methanogen genomes added on 16^th^ of December 2025 were not included in statistical analyses because of their low TR densities (Supplementary Table 19). Annotation and proteome files were retrieved from NCBI when available. Phylogenetic trees were constructed from 76 archaeal single-copy marker genes using IQ-TREE v2.3.6 with the ‘-m MFP −B 1000’ flags as implemented in GToTree v1.8.6^58^ and visualized in iTOL v7.5.1^59^.

### Tandem repeat identification and characterization

Tandem repeats were identified at both the nucleotide and amino acid levels using a custom Python script, which builds on MUMmer v3.23^60^ as previously described (https://github.com/rohansachdeva/tandem_repeats)^25^. Stringent thresholds were used allowing no mismatch and detecting only nucleotide TRs of a total region length ≥ 50 nt and ≥ 3 repeat units; aaTRs were searched in the predicted proteomes using thresholds of ≥ 16 aa region length and ≥ 3 TR units. These criteria are more conservative than those commonly applied in bacterial microsatellite surveys and restrict the analysis to perfect, slippage-prone tracts. To evaluate how many imperfect TRs were excluded, a modified version of the script allowing up to two mismatches per repeat unit was applied (Supplementary Note 1). Degenerated protein repeats in *Methanosarcina* and *M. ruminantium* (NC_013790.1) proteomes were detected with lowered thresholds (≥ 9 aa length, ≥ 3 TR units) and repetitive motifs (≥ 50 nt and ≥ 3 repeat units) within *Methanosarcina* disaggregatase-encoding genes were identified using Geneious Prime 2026.1.1 (https://www.geneious.com; also used for manual sequence inspection described below; Supplementary Table 13).

When genome annotation files were available, each TR was assigned to one of three genomic categories based on overlap with ORF coordinates (Biopython): (i) ‘genic’, (ii) ‘ORF overlapping’, and (iii) ‘intergenic’. Proportions were computed per genome and then averaged across genomes to avoid the summary being dominated by the most TR-rich genomes and were compared against the coding density of each genome, defined as the ratio of the non-redundant coding union to the total assembly length. Because distributions of TR length, unit length and unit number are strongly right skewed, these are reported as medians (Supplementary Fig. 1).

Identical TRs across genomes were identified by global pairwise alignment (Biopython’s PairwiseAligner) of the extracted TR unit sequences and their reverse complements, accounting for all possible circular rotations of the repeat unit (Supplementary Table 4.2) and requiring 100% identity.

### Statistical analysis of TR density, composition and genomic context

To identify drivers of TR density in 170 methanogen genomes, we fitted a multivariable Negative Binomial Generalized Linear Mixed-Effects Model on raw TR counts using glmmTMB implemented in R with log-transformed genome size, broad ecosystem classification, and the predicted parameters mentioned above as fixed effects and genus identity as a random intercept. Fixed effect significance was evaluated using Type II Wald Chi-squared (*χ*^2^) tests and the random intercept using a Likelihood Ratio Test: parameter stability across habitats with unequal sample sizes (n = 8–34) was confirmed by stratified bootstrapping (Supplementary Note 1). The enrichment of TRs in *Methanosarcina* genomes was further tested using a Wilcoxon rank-sum test.

The compositional bias of nucleotide TRs and non-TR sequences was assessed as described previously^25^: genomes were divided into TR and non-TR segments into 50 nt long substrings, and nucleotide frequencies aggregated by category were compared using the Kruskal-Wallis test with FDR correction according to the Benjamini-Yekutieli method^61^.

To assess the genomic context of intergenic TRs, the 28 dereplicated (99% ANI) *Methanosarcina* genomes with predicted coding sequences were used. ORFs were characterized using KofamScan v1.3.0^62^ against the KEGG Orthology (KO) database (ko00001; accessed March 10, 2026) retaining the top-scoring hit per ORF (*E*-value < 0.001) and the direct genomic neighborhood of each TR was defined as its immediate upstream and downstream flanking genes. Enrichment or depletion of functional groups and their higher hierarchical categories (KO level C) within these neighborhoods was tested with an empirical permutation test (1,000 iterations) comparing the observed gene counts within the TR neighborhoods against a null distribution of equally sized random samples from the annotated background. Deviations were quantified as permutation-derived *Z*-scores with two-sided empirical *p*-values and Benjamini-Hochberg (BH) FDR correction^63^, and functional categories with fewer than 10 observed hits or only one KO were excluded.

### Statistical characterization of TR-containing RNAs, amino acid TRs, and TR-proteins

Potential binding target mRNAs of the TR-containing RNAs were predicted with the IntaRNA web server (v3.4.1, standard parameters) against the *M. mazei* Gö1 reference genome (NC_003901.1). TR-proteins in *Methanosarcina* were characterized in the proteomes of the same 28 dereplicated genomes. Subcellular localization was predicted with PSORTb v3.0^64^ in archaeal mode; proteins with an ‘unknown’ localization assignment (14.0% of the global proteomic background and 28.8% of TR-proteins) were excluded from downstream sub-cellular enrichment analyses (Supplementary Note 1). Amino acid sequences of TR-proteins were profiled using KofamScan v1.3.0^62^, IDRs were predicted with MobiDBLite v2.0^65^, and functional annotations were validated or curated using InterProScan v5.69-101.0^66^.

For each unique TR-protein, the amino acid coordinates of functional domains and IDRs were mapped against TR boundaries, and domains were then grouped as TR-driven (the TR makes up at least 50% of the domain), partial overlap (less than 50%), or separate elements encoded on the same protein. To identify domains preferentially located at the cell surface that co-occur with aaTRs, each unique InterPro domain was tested for non-random association with predicted cell-surface localization (PSORTb categories ‘extracellular’ and ‘cell wall’), using two-sided Fisher’s exact tests with sample odds ratios and BH-adjusted *p*-values, and surface-enriched domains were intersected with the set of aaTR-containing proteins (Supplementary Table 12). Proteins harboring multiple TRs were manually inspected in Geneious Prime and categorized as a single continuous TR region (proximal TRs with minimal sequence divergence) or multiple independent TR regions (spatially separated or exhibiting distinct consensus sequences; Extended Data).

Enrichment or depletion of KO level C categories among TR-proteins was tested with the permutation approach described above (5,000 iterations), and higher functional categories completely absent from the TR-protein pool were profiled by their absolute copy number and number of individual KOs (filtered for n > 3) within the global background proteome.

Proteins of the 28 dereplicated proteomes were clustered using MMSeqs2 v14-7e284^67^ (--min-seq-id 0.50 -c 0.7 --cluster-reassign). The functional prediction of each cluster was based on the most abundant prediction within the cluster, and where this majority-vote annotation conflicts with the InterPro domain content of the cluster, both are reported (Extended Data Table 2). Identical aaTR sequences across different proteins were identified after standardizing motifs to a single canonical rotation. Differences between TR-proteins and the proteomic background, and among subcellular localizations, were tested using Fisher’s exact tests (localization, multi-TR prevalence, IDR and domain enrichment), Kruskal-Wallis tests with post-hoc Dunn’s tests (protein length, TR coverage), Chi-squared (*χ*^2^) tests of independence (amino acid composition across localizations) and paired Wilcoxon signed-rank tests (amino acid composition relative to the background); full details are given in Supplementary Note 1.

### Whole genome sequencing (ONT)

Cells of *M. mazei* DSMZ and Ref_ETH cultures were harvested during exponential growth by centrifugation at 8,000 g for 5 min and genomic DNA was isolated using Qiagen’s MagAttract HMW DNA Kit. *M. mazei* Ref_WT cells were harvested by centrifugation at 13,000 g for 2 min and genomic DNA was isolated using the Wizard Genomic DNA Purification Kit (Promega) following the manufacturers’ protocols.

Genomic DNA was purified using AMPure XP beads, libraries were prepared using the Native Barcoding Kit V14 (SQK-NBD114.24) and sequenced on the Oxford Nanopore PromethION 2 Solo system (read yields in Supplementary Note 1). Reads were basecalled using the model dna_r10.4.1_e8.2_400bps_sup@v5.0.0 with Dorado v7.8.3, filtered for Q-score ≥ 15 and ≥ 1,000 bp using chopper v0.10.0^68^ and subsampled with Filtlong (https://github.com/rrwick/filtlong; --target_bases 1Gb, 400Mb or 210Mb depending on read quality and sequencing depth). Genomes were assembled using the reference-based assembler Rebaler v0.2.0 (https://github.com/rrwick/Rebaler) using NC_003901.1. TR regions were identified in the assembled genomes as described above and ORFs were transferred from the reference genome to the assemblies using liftoff (https://github.com/agshumate/Liftoff).

### Evolution of TRs

TR similarity between the different *M. mazei* assemblies was analyzed using global pairwise alignments as described above (Supplementary Table 5.1 and 5.2). In-sample TR variance was assessed using tandem-genotypes v1.9.2^69^. Quality-filtered reads were randomly subsampled to the same number of bases across samples using seqkit v0.10.1^70^ and mapped to the assembled genomes using LAST^71^ with the recommended parameters for long reads (‘-uRY4’). To assess the rate and variant frequency of single nucleotide variants (SNVs), the same set of reads was mapped against the assembled genomes using minimap2 v2.22^72^ with the parameters ‘-x lr:hq --secondary=no’ and screened for SNVs using Clair3 v2.0.2^73^ with the ONT specific model.

### Expression of TRs

Ribosome profiling (Ribo-seq) and RNA sequencing (RNA-seq) datasets were downloaded from the NCBI Sequence Read Archive (SRA) on 13 July 2026 under BioProject accession PRJNA1004312^7^ (GEO series GSE240615). Adapter sequences were trimmed and low-quality reads were removed using Trimmomatic v0.39^74^ in single-end mode, ribosomal RNA reads were filtered using Bowtie2 v2.5.4^75^ (‘--very-sensitive-local’) mapped against the SortMeRNA database v4.3^76^, and the remaining reads were aligned to the reference genome (NC_003901.1) using Bowtie2 with the same parameters (Supplementary Note 1); genome coverage was analyzed using Geneious Prime. For each TR region the expression was categorized into “low” (5–29 reads), “moderate” (30–100 reads), and “high” (> 100 reads).

For validation of TR expression using long-read sequencing, *M. mazei* Gö1 cultures were harvested during exponential growth (OD_600_ ∼ 0.2) by centrifugation at 13,000 g for 2 min, total RNA was extracted using RotiZol and DNaseI treated as described previously^77^. Ribosomal RNA was depleted following siTOOLs Pan-Archaea riboPOOL kit (dp-K012-27) with subsequent ethanol precipitation. rRNA depleted RNA samples were polyA tailed following the Oxford Nanopore protocol. Libraries were prepared using the cDNA-PCR Barcoding Kit V14 (SQK-PCB114.24) and sequenced using the Oxford Nanopore PromethION 2 Solo system (Supplementary Note 1). After base calling using the model dna_r10.4.1_e8.2_400bps_sup@v5.0.0 with Dorado v7.9.8, raw reads were length filtered for ≥ 15 bp using chopper v0.10.0^68^, artificially added 3’ polyA tails were removed using cutadapt v4.4^78^ with following parameter: “-a A{10} -e 3 -j 0”. Reads were quality filtered for Q-score ≥ 10 before remaining rRNA reads were removed using minimap2 v2.22^72^ (“-ax splice -uf”) with the sensitive SortMeRNA^76^ database together with the known rRNA sequences of the *M. mazei* Gö1 (NC_003901.1) genome. Non ribosomal reads were extracted using samtools v1.17^79^, mapped to the assembled genome using minimap2 v2.22 with the same parameters and analyzed in Geneious Prime.

### Proteomics data analyses

All vendor-format raw mass spectrometry files from *M. mazei* Gö1 (PXD004325; 72 files), *M. acetivorans* C2A (PXD022016; 15 files), and *Methanobrevibacter olleyae* YLM1 (PXD070174; 10 files) were retrieved from the PRIDE database^80^ and reanalyzed with the same search settings for all three datasets in Proteome Discoverer 3.3 (v3.3.1.110) using CHIMERYS^81^ with organism-specific whole-proteome protein databases together with a common-contaminant sequence database^82^ (Supplementary Note 1).). One PXD022016 file (181207_M42_rep3) could not be searched by CHIMERYS because one of its spectra contains no peaks, so identifications for this dataset derive from 14 of its 15 files. Peptide-spectrum matches (PSMs) and peptides with global q-values below 0.01 (1% FDR) were retained, proteins supported by at least one retained peptide were grouped using strict parsimony and evaluated at a protein-level FDR threshold of 1%, and only proteins with q-values below 0.01 were counted as detected proteins and used to calculate detection rates.

### TR evolution of the ΨM2 tape measure protein

The ΨM2 reference genome (NC_001902.1) was downloaded from NCBI on 2 May 2025. Genes were grouped in functional categories using PhageScope^83^ and the TR region of the reference genome was verified as described above. Genomic ONT long reads were generated and quality filtered as part of a previous study^37^ (SRA accessions SRX35246099–SRX35246103). Reads from all five samples were pooled, and TR variation was analyzed as described above without subsampling. Tape measure protein (IPR010090; AAC27060.1 in AF065411) sequences were manually edited to generate variants representing low (2 units), mean (13 units), and high (21 units) repeat unit counts alongside the reference sequence (10 units), and protein domains were predicted using InterProScan^66^ in Geneious Prime 2026.1.1.

### Methanogen plasmids and viruses

Genomes of methanogen plasmids were retrieved from NCBI (accession numbers are provided in Supplementary Table 16), repeats were identified using the workflow described above, and GC-skew was calculated using the Python script integrated within iRep^84^. Viral genome sequences targeting methanogen hosts were compiled from multiple public databases and literature-derived resources (Supplementary Table 20), generating an initial dataset of 447,591 viral genomes, which were dereplicated using seqkit v2.12.0^85^ (rmdup -s) to 407,171 non-redundant genomes and screened for tandem repeats as described above. GeNomad v1.8.1^86^ was used to confirm that all sequences were classified as viral and to assign viral taxonomy. Protein-coding genes were predicted using Prodigal-gv v2.11.0^87^, and genome annotation was performed using Pharokka v1.9.1^88^, eggNOG-mapper v2.1.12^89^, KofamScan v1.3.0^62^, InterProScan v5.69-101.0^66^, and Colabfold v1.5.5^90^ together with FoldSeek v8-ef4e960^91^ (against PDB released January 1, 2024) to improve functional characterization of viral proteins using a cutoff e-value of 10^-4^. Annotations were manually assessed and grouped into functional categories (Supplementary Table 18). Putative host associations were inferred using iPHoP v1.4.2^92^, from which viruses predicted to infect methanogenic archaea were selected for downstream analysis. Genome quality was evaluated using CheckV v1.0.3^93^, retaining genomes with ≥50% completeness and at least two predicted genes. TR loci were categorized according to their genomic position relative to predicted ORFs.

### Statistics and reproducibility

Statistical analyses were performed in R and Python. All tests were two-sided, and *P* values from multiple comparisons were adjusted using the Benjamini-Hochberg procedure unless stated otherwise. Read-level TR heterogeneity was quantified in three independently propagated *M. mazei* Gö1 cultures. No statistical method was used to predetermine sample sizes, and randomization and blinding were not applicable to this study.

## Data availability

Oxford Nanopore whole-genome and transcriptome sequencing data generated in this study are available under NCBI BioProject PRJNA1548339. The annotated *M. mazei* Gö1 genome with mapped TR regions, the three *M. mazei* Gö1 assemblies, together with the tandem repeat gff/bed files, alignment and tree files, protein clustering, disorder and domain annotations, and full IntaRNA and PSORTb output, are available on Zenodo (https://doi.org/10.5281/zenodo.23493132). All other data are publicly available: genome, plasmid and viral genome accessions are listed in Supplementary Tables 1, 2.1, 3, 16, 19 and 20; RNA-seq and Ribo-seq data under PRJNA1004312 (SRR25604690– SRR25604697); ΨM2 reads under PRJNA1529606 (SRX35246099–SRX35246103); proteomics raw files under PRIDE PXD004325, PXD022016, PXD070174, with the CHIMERYS reanalysis available on Zenodo (https://doi.org/10.5281/zenodo.23493132).

## Code availability

Custom scripts for tandem repeat detection, including the mismatch-tolerant modification, enrichment testing and statistical analysis are available at https://github.com/envmicrobiol-ethz/Haiss_et_al_2026 and archived at Zenodo (https://doi.org/10.5281/zenodo.23493132). The original repeat detection script is available at https://github.com/rohansachdeva/tandem_repeats.

## Author Contributions

The study was conceived and designed by SH and MCS. SH performed TR analyses on publicly available genomes and proteomes, constructed phylogenetic trees, and carried out comparative genomics, enrichment analyses, RNA-seq and Ribo-seq data analyses with input from MAT, and sRNA target predictions. Genomic DNA of *M. mazei* Gö1 was extracted by KW and SH. Total RNA was extracted by KW with rRNA depletion carried out by SH. SH and JH performed library preparation, ONT whole-genome sequencing, and transcriptome sequencing. SH performed SNV calling for the *M. mazei* Gö1 and read-level variant analyses for the *M. mazei* Gö1 and ΨM2 datasets. JH analyzed the ΨM2 whole-genome sequencing data; SH, JH, and MCS interpreted these data together. CD analyzed publicly available proteomics datasets, with input from SH. MCS and SH analyzed TRs in methanogen plasmids. EJS generated and analyzed methanogen virus datasets identified TRs in predicted methanogen-associated viruses and interpreted these data together with SH. SS and BM performed experiments with *M. marburgensis* and the ΨM2 virus. KW and RAS provided strains and KW, RAS, and MAT advised on *M. mazei*. SH generated the figures, with contributions from EJS and JH. SH and MCS wrote the manuscript with input from all authors. Funding was acquired by MCS.

## Acknowledgements

Funding for this research was provided by the SNSF Starting Grant (SNF 226487). We thank Donat Crippa, Bledina Dede, Taylor Priest, Grayson Chadwick for scientific input, and Lukas Wanner for technical support. We thank the Genetic Diversity Centre (GDC), ETH Zurich, for providing facility access and technical support.

