## Extended Data for "Tandem repeats in methanogenic archaea and their viruses reveal a shared potential for rapid genomic diversification"

This Extended Data file contains results, figures and tables supporting the manuscript's primary findings. Additional supporting data can be found in the Supplementary Information file and Supplementary Tables 1–20.

### Results

#### Many TRs are transcribed and show ribosome occupancy

Twelve TRs showed significant higher read coverage than their immediate genomic neighborhood, which may reflect genuinely elevated local transcription (4 instances), ribosome occupancy (7 instances), or both (1 instance). This could arise if the TR forms part of an independently transcribed small RNA, or inflation because the sequenced culture carries more units at that locus than the reference.

#### Genic TRs encode disordered extensions concentrated in cell-surface proteins

In *Methanosarcina*, 110/960 TR-proteins harbored multiple aaTRs. In most such cases (n=75), these aaTRs differed only slightly in sequence and were located in close proximity, suggesting local within-region evolution. Conversely, 31.8% of multi-TR proteins possessed distinct aaTR regions. Proteins with unknown cellular localization displayed a markedly higher TR coverage, measured as the proportion of total protein length comprising TRs, than those localized to the cytoplasm or membrane ($p < 0.002$).

To assess if sequence-homologous proteins are enriched in aaTRs, we performed protein family clustering (50% identity, 70% coverage), which placed 90.7% of all 95,121 proteins into 7,228 clusters, leaving 8,885 singletons (Supplementary Table 20). The clusters richest in aaTRs were dominated by surface proteins (Extended Data Extended Data **Table *2***). The top two, cluster_05737 (10/11 proteins) and cluster_10043 (10/13), are extracellular or cell wall-associated, carrying signal peptides, IDRs and PKD domains (IPR035986) or cell-adhesion related domains (CARD; IPR011635), respectively. Cluster_05737 additionally carries a heme domain (IPR011048) and, in two homologs, an MCP signaling domain embedded in the TR region. Since *Methanosarcina* does not denitrify, we propose the heme acts as a ligand- or oxygen-sensing site[94,95] rather than supporting the nitrous oxide reductase annotation (IPR011045). Three further clusters comprise IDR-containing proteins of unknown function and one a secreted carboxypeptidase, leaving only cluster_13854 as clearly cytoplasmic.

#### TRs occur in viruses that infect methanogens

No TRs were detected in MetSV, infecting *Methanosarcina*[10]; Blf4 infecting *Methanoculleus*[96]; Drs3 infecting *Methanobacterium*[97]; MFTV1 infecting *Methanocaldococcus*[98]; or MSTV1 infecting *Methanobrevibacter*[99]. Quality filtered (≥50% CheckV completeness) TR-containing vMAGs were predominantly classified as unclassified Caudoviricetes (85.5%), and were predicted to infect *Methanobrevibacter* A (~57%), *Methanosarcina* (17%), *Methanofollis* (~9%), and *Methanothermobacter* hosts (4%).

### Extended Figures

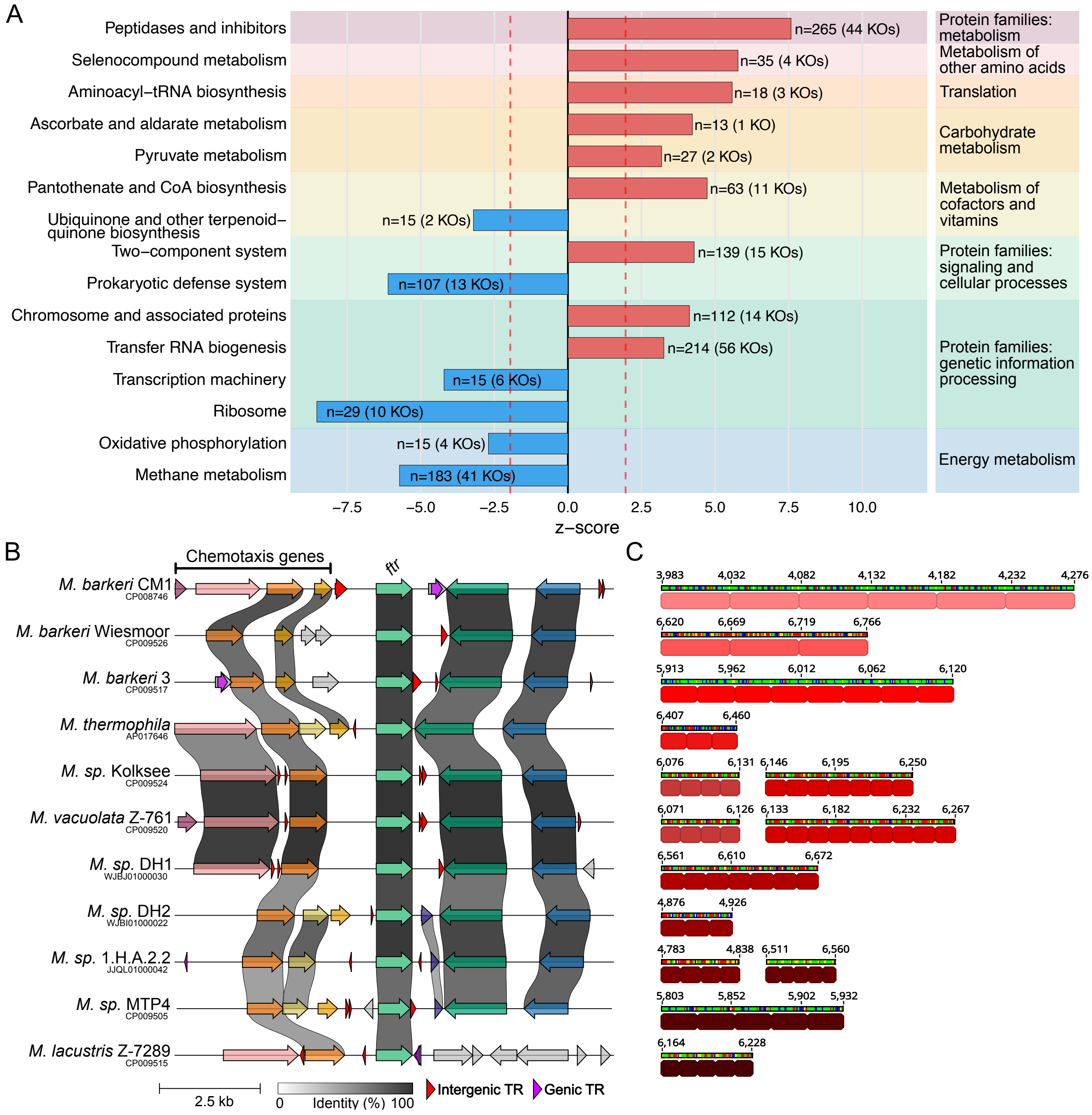


**Extended Data Fig. 1 | Intergenic TRs in *Methanosarcina* are associated with specific functional gene categories. (A)** Enrichment analysis of functional gene categories in direct TR neighborhoods. Bars represent *Z*-scores (positive: enrichment; negative: depletion; dashed red lines: significance threshold at ± 1.96, $p < 0.05$). Only categories with > 10 TR-adjacent genes are shown. Numbers indicating total associated genes and unique KEGG Orthology (KO) groups are shown in parentheses. Background shading highlights higher-order functional hierarchies. **(B)** Gene cluster alignment of the formylmethanofuran-tetrahydromethanopterin N-formyltransferase (*ftr*) locus across *Methanosarcina* genomes, illustrating frequent but positionally heterogeneous TR associations. The *ftr* gene is flanked by upstream chemotaxis-related genes and downstream TraB family and CBS domain-containing proteins. **(C)** TRs proximal to *ftr* show extensive variation in unit length, sequence (red color gradient), and unit number.

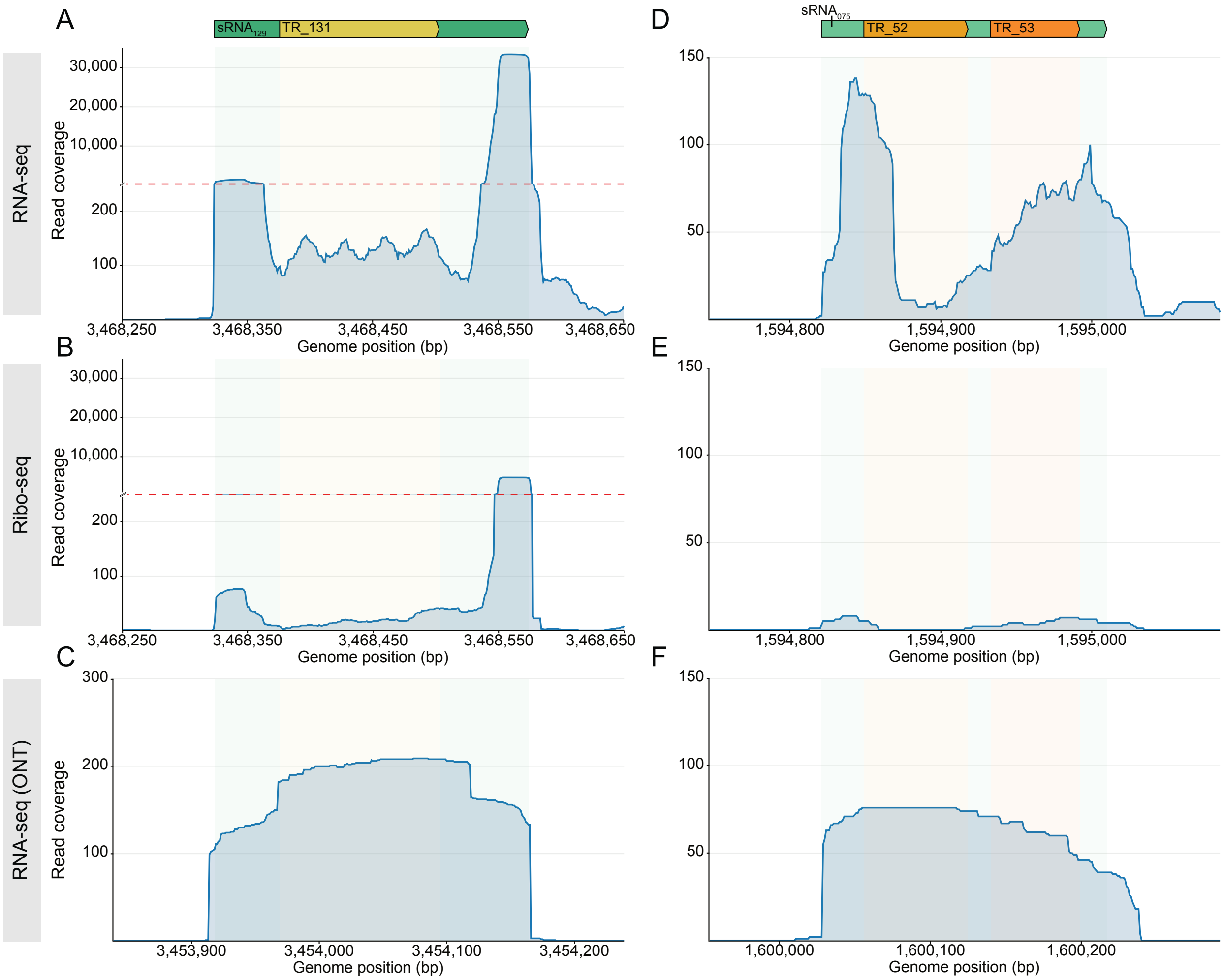


**Extended Data Fig. 2 | Expression profiles of TRs in *M. mazei* Gö1. (A–C)** Read coverages for TR_131, located within verified sRNA129. **(D–F)** Read coverages for TR_52 and TR_53, located within verified sRNA75. Panels depict short-read RNA-seq **(A, D)** and Ribo-seq **(B, E)** data re-analyzed from Tufail *et al.* (2024)[7], alongside long-read Oxford Nanopore Technologies (ONT) sequencing data **(C, F).**

### Extended Tables

Complete datasets are available as separate CSV files (Extended Data Tables 1 and 2), with key findings summarized in the abbreviated tables below.

**Extended Data Table 1 | Functional genes most frequently associated with proximal TRs in *Methanosarcina*.** The TR rate is the percentage of genes within each KO group located proximal to a TR; counts (adjacent/total) are in parentheses. The mean background TR association rate across all genes is 7.26%, representing the expected frequency under stochastic genomic distribution. All listed associations are significantly enriched above this background ($p < 0.001$).

| KEGG Orthology | TR rate | *p* value | # Genomes |
|---|---|---|---|
| **K00096:** glycerol-1-phosphate dehydrogenase [NAD(P)$^+$] [EC:1.1.1.261] | 42.86% (12/28) | $2.15 \times 10^{-7}$ | 12 |

| | | | |
|---|---|---|---|
| **K18933:** tyrosine decarboxylase / aspartate 1-decarboxylase [EC:4.1.1.25, 4.1.1.11] | 42.86% (12/28) | $2.15 \times 10^{-7}$ | 12 |
| **K07574:** RNA-binding protein | 40.74% (11/27) | $1.28 \times 10^{-6}$ | 11 |
| **K07765:** S2P endopeptidase [EC:3.4.24.85] | 40.74% (11/27) | $1.28 \times 10^{-6}$ | 11 |
| **K00672:** formylmethanofuran—tetrahydro-methanopterin N-formyltransferase [EC:2.3.1.101] | 37.93% (11/27) | $2.96 \times 10^{-6}$ | 11 |
| **K08963:** methylthioribose-1-phosphate isomerase [EC:5.3.1.23] | 35.71% (10/28) | $1.57 \times 10^{-5}$ | 10 |
| **K10725:** archaeal cell division control protein 6 | 29.51% (18/61) | $1.94 \times 10^{-7}$ | 16 |
| **K22373:** lactate racemase [EC:5.1.2.1] | 25.0% (15/60) | $1.86 \times 10^{-5}$ | 8 |
| **K00549:** 5-methyltetrahydropteroyltriglutamate--homocysteine methyltransferase [EC:2.1.1.14] | 24.29% (17/70) | $8.06 \times 10^{-6}$ | 11 |
| **K01953:** asparagine synthase (glutamine-hydrolysing) [EC:6.3.5.4] | 24.0% (12/50) | $1.90 \times 10^{-4}$ | 12 |

**Extended Data Table 2 | *Methanosarcina* protein clusters with the highest proportion of TR-proteins.** The frequency of TRs per cluster is expressed as the aaTR rate. The individual aaTR unit sequences are listed together with the corresponding protein numbers. Predicted cellular localizations, KEGG Orthology (KO) identifiers, and functional annotations are listed for each cluster.

| Cluster | aaTR rate | Cellular localization | [aaTR units] and frequency | cluster_KOs | Functional annotation (InterPro) |
|---|---|---|---|---|---|
| 05737 | 90.91% (10/11) | cellwall [3]; extracellular [1]; unknown [7] | [AETETEPLEEPLEEPTETETEPSEP]$_{1}$; [DNITIPE]$_{4}$; [DNVTIPE]$_{2}$; [EENITVP]$_{1}$; [EENVTVP]$_{1}$; [EEVTEPT]$_{2}$; [EEVTPT]$_{4}$; [EEVVEPT]$_{1}$ | K20932 [11] | methyl-accepting chemotaxis protein; cell adhesion |
| 10043 | 76.92% (10/13) | extracellular [8]; unknown [5] | [DDKPSDKPG]$_{1}$; [DDTDNGKNVT]$_{1}$; [DDTGDDTS]$_{1}$; [DDTS]$_{1}$; [DNGSDNSS]$_{1}$; [DNGTS]$_{3}$; [DNNTGDNSS]$_{1}$; [DNTSGNTT]$_{1}$; [DTNVTI]$_{3}$; [DTNVTN]$_{2}$ | K04079 [1]; K09637 [2]; K13018 [1] | cell adhesion; transmembrane proteinase |
| 10508 | 54.55% (6/11) | cytoplasmic [4]; unknown [7] | [AAENVTEDV]$_{1}$; [ANVTDD]$_{3;}$ [DDNMS]$_{2}$ | - | unknown |
| 00013 | 50.0% (5/10) | cytoplasmic [2]; unknown [8] | [AADEE]$_{1}$; [AADNMTVEDNDS]$_{1}$; [AEEEV]$_{2}$; [DENVT]$_{1}$ | K25373 [1] | unknown |
| 07705 | 48.0% (12/25) | cytoplasmic [25] | [AEEE]$_{2}$; [AEEEAEEEE]$_{1}$; [AEEEE]$_{2}$; [AEESEEESEEE]$_{1}$; [AEPEEE]$_{1}$; [AEVEEE]$_{1}$; [ALEEEEE]$_{1}$; [DLEE]$_{2}$; [EEESSL]$_{1}$; [EEEVQP]$_{1}$; [EEKPEI]$_{1}$ | K11600 [25] | exosome complex component RRP41 |
| 13854 | 46.67% (7/15) | cytoplasmic [14]; unknown [1] | [DLTE]$_{1}$; [EEITEEVT]$_{2}$; [EEVT]$_{1}$; [LNNS]$_{1}$; [MNNS]$_{2}$ | K20333 [15] | toxoflavin biosynthesis protein ToxD; sulfatase-modifying factor enzyme |
| 13396 | 35.71% (10/28) | cytoplasmic [27]; unknown [1] | [ASNIPD]$_{2}$; [DSIPDSIS]$_{1}$; [DSIS]$_{2}$; [DSVH]$_{3}$; [DSVS]$_{3}$; [DTPNTL]$_{1}$; [ESSV]$_{1}$ | K02323 [28] | DNA polymerase II small subunit [EC:2.7.7.7] |

| | | | | | |
|---|---|---|---|---|---|
| 16108 | 30.0% (3/10) | extracellular [7]; unknown [3] | [ETSQTLT]$_3$ | K05999 [10] | xanthomonalisin [EC:3.4.21.101] |
| 08872 | 30.0% (3/10) | unknown [10] | [DPVPTDTP]$_3$ | K14201 [1] | unknown |
| 00658 | 28.57% (4/14) | cytoplasmic [14] | [CNISVSDDK]$_1$; [CNPDD]$_1$; [DISDK]$_1$; [DKIIP]$_1$; [DKSIS]$_1$; [EQQETL]$_1$ | K03407 [14] | two-component system, chemotaxis family, sensor kinase CheA [EC:2.7.13.3] |
| 11050 | 24.0% (6/25) | cytoplasmic [25] | [DSKIQ]$_2$; [EEGG]$_1$; [EEGKEKK]$_1$; [EEREKGEKI]$_1$; [EERKEKG]$_1$; [EKEV]$_1$ | K03407 [25] | two-component system, chemotaxis family, sensor kinase CheA [EC:2.7.13.3] |
| 00543 | 27.8% (5/18) | extracellular [11]; unknown [7] | [DDQSGF]$_2$; [DNQTESL]$_1$; [ENQMEVNES]$_1$; [ESEYEY]$_1$ | K07218 [18] | nitrous oxidase accessory protein nosD |
