## Supplementary Information for "Tandem repeats in methanogenic archaea and their viruses reveal a shared potential for rapid genomic diversification"

This Supplementary Information file contains Supplementary Figures, a list of Supplementary Tables supporting the manuscript's findings and Supplementary Note 1 with Extended Methods. Supplementary Tables are provided as separate CSV files.

### Supplementary Figures:

**Supplementary Fig. 1 | Structural properties of nucleotide TRs in 87 methanogen genomes.** Frequency of TR region lengths **(A)**, repeat unit lengths **(B)**, repeat unit counts per repeat **(C)** and repeat GC content **(D)**. ........ 3

**Supplementary Fig. 2 | Nucleotide composition of TR regions compared to their respective genomes.** Each data point displays the TR-versus-background variation calculated for **(A)** 87 methanogen genomes and **(B)** 28 Methanosarcina genomes. TR regions are significantly enriched in adenine and thymine, and depleted of guanine and cytosine (***, p < 0.001). ........ 4

**Supplementary Fig. 3 | TR sequences diverge faster than core genomic backgrounds. (A)** Heatmap showing the percentage of shared TR sequences between a selection of methanogen genomes. **(B)** Proportions of shared TRs plotted against average nucleotide identity (ANI) from pairwise comparisons of Methanosarcina genomes. The red dotted line indicates the 95% ANI species boundary, and orange data points highlight intra-species comparisons within M. mazei. Inset: Number of TR regions in M. mazei Gö1 (orange) compared to closely related genomes. **(C)** An expanded view of the data in panel B, demonstrating that TR divergence occurs even between closely related strains (ANI > 99.5%; red dotted line). **(D)** Total TR region counts in closely related strains (blue) relative to M. mazei Gö1 (orange). ........ 5

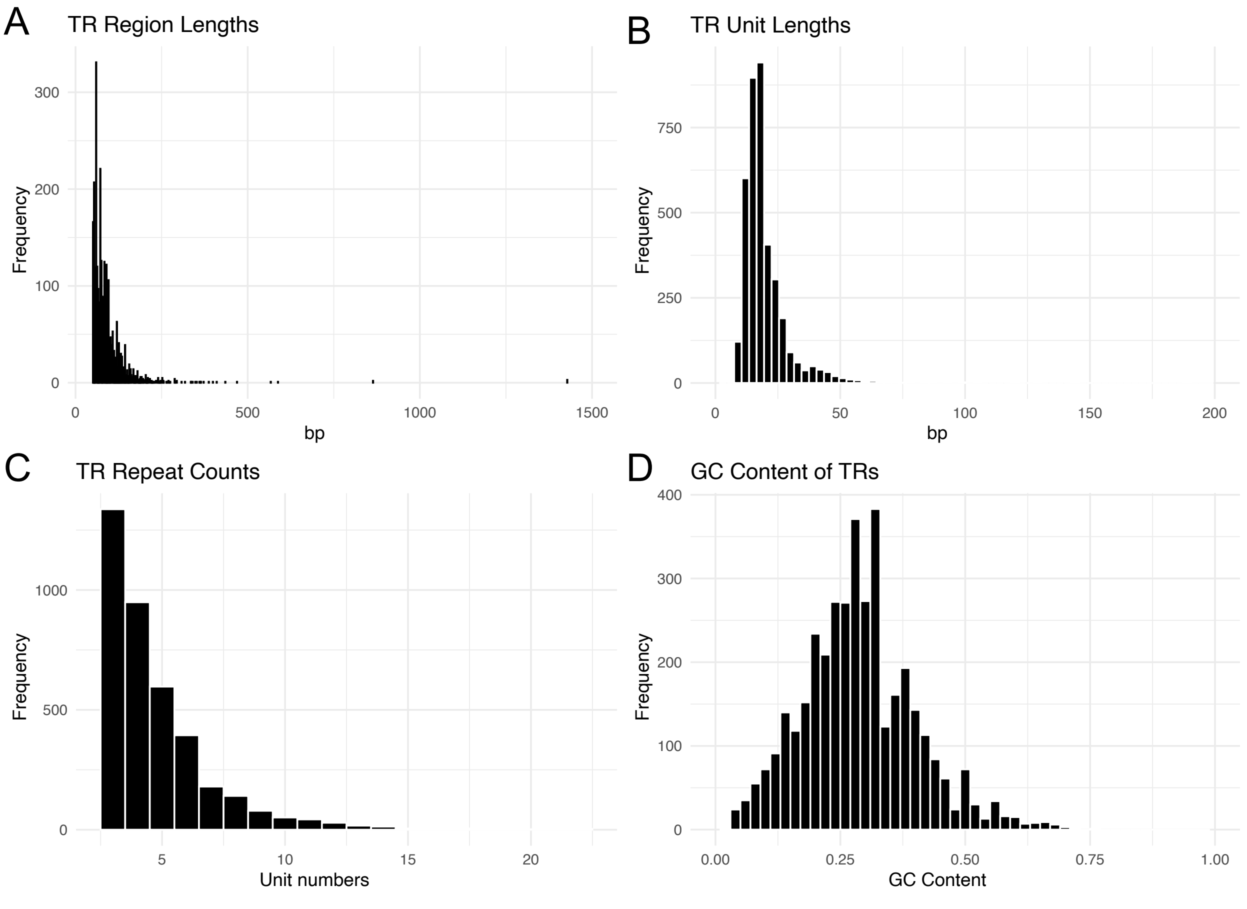

**Supplementary Fig. 1 | Structural properties of nucleotide TRs in 87 methanogen genomes.** Frequency of TR region lengths **(A)**, repeat unit lengths **(B)**, repeat unit counts per repeat **(C)** and repeat GC content **(D)**.

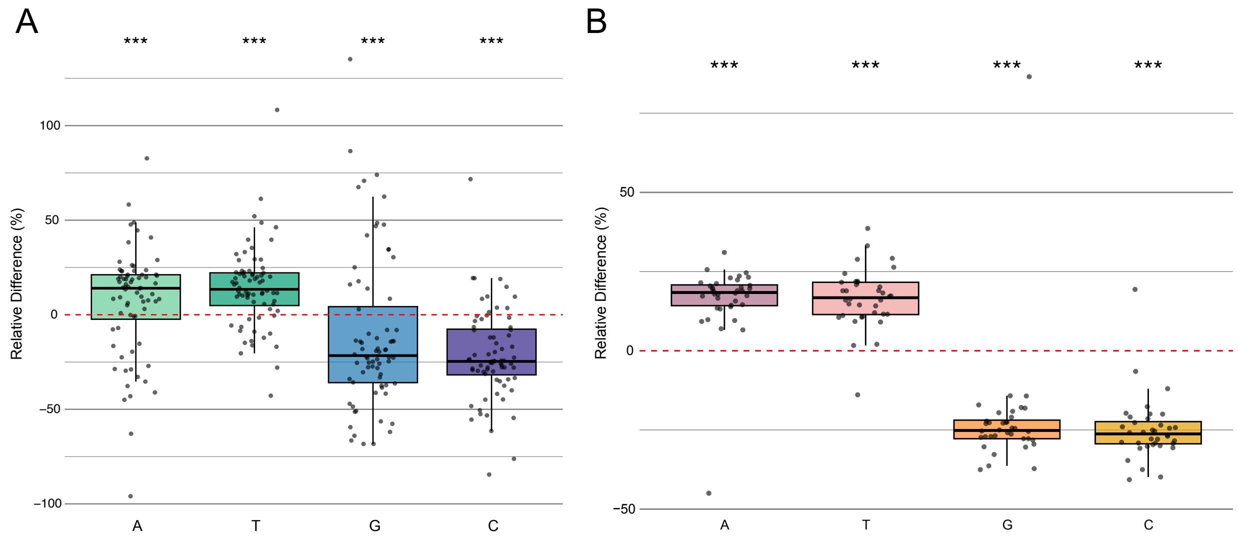

**Supplementary Fig. 2 | Nucleotide composition of TR regions compared to their respective genomes.** Each data point displays the TR-versus-background variation calculated for **(A)** 87 methanogen genomes and **(B)** 28 *Methanosarcina* genomes. TR regions are significantly enriched in adenine and thymine, and depleted of guanine and cytosine (***, $p < 0.001$).

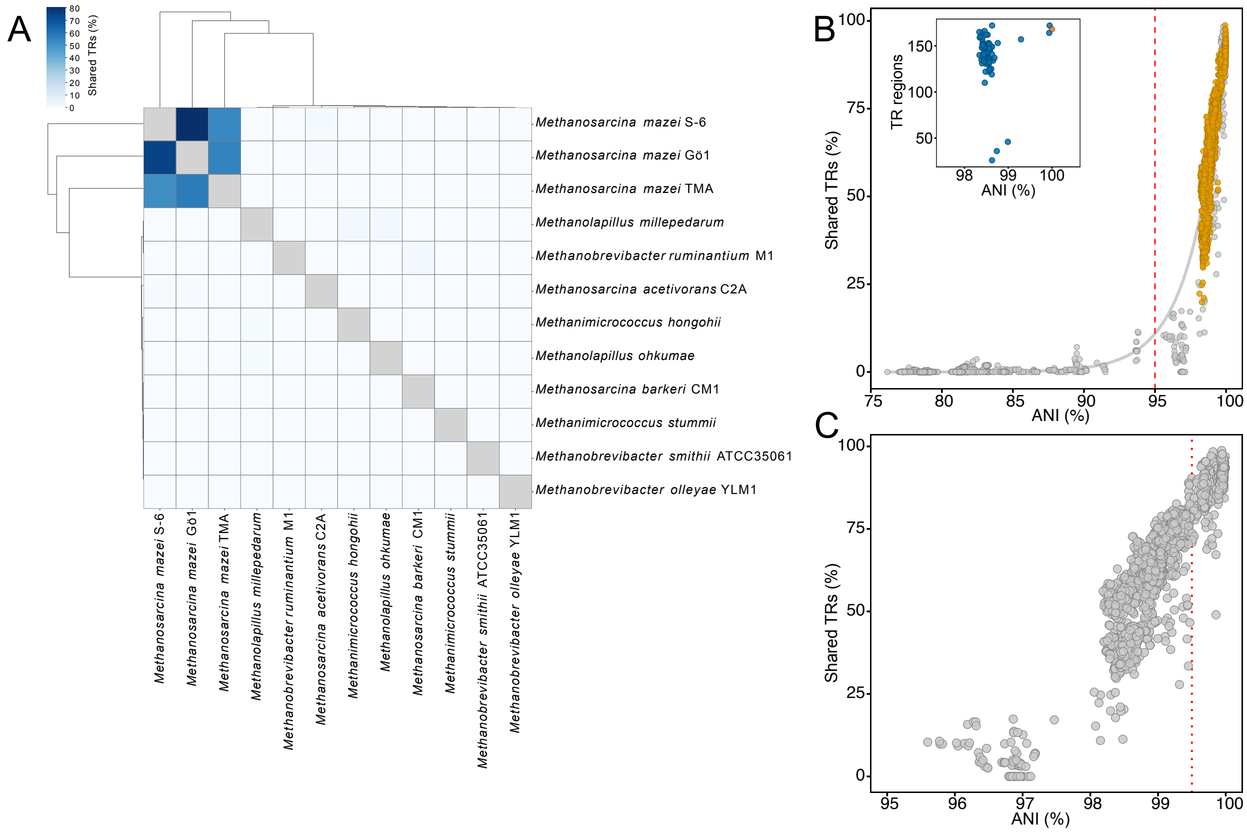

**Supplementary Fig. 3 | TR sequences diverge faster than core genomic backgrounds. (A)** Heatmap showing the percentage of shared TR sequences between a selection of methanogen genomes. **(B)** Proportions of shared TRs plotted against average nucleotide identity (ANI) from pairwise comparisons of *Methanosarcina* genomes. The red dotted line indicates the 95% ANI species boundary, and orange data points highlight intra-species comparisons within *M. mazei*. Inset: Number of TR regions in *M. mazei* Gö1 (orange) compared to closely related genomes. **(C)** An expanded view of the data in panel B, demonstrating that TR divergence occurs even between closely related strains (ANI > 99.5%; red dotted line). **(D)** Total TR region counts in closely related strains (blue) relative to *M. mazei* Gö1 (orange).

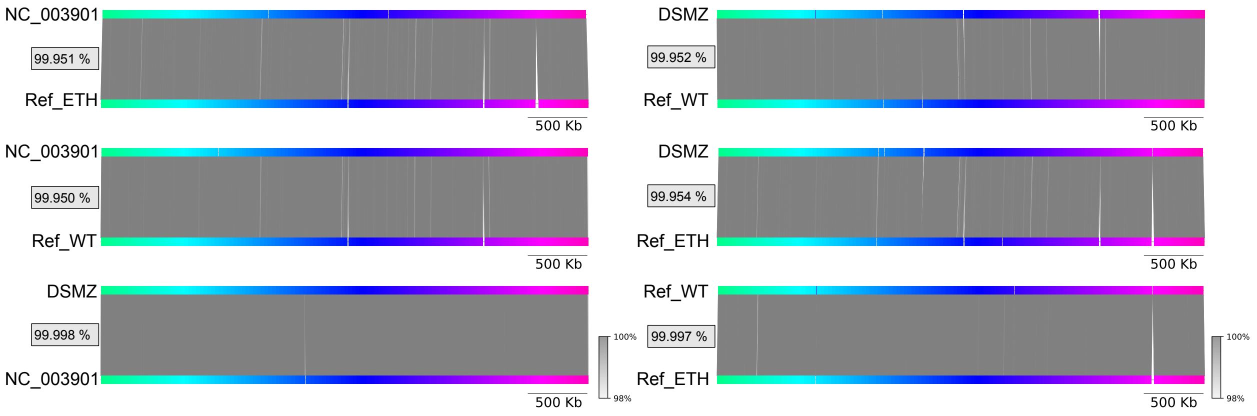

**Supplementary Fig. 4 | Pairwise ANI of *M. mazei* Gö1 cultures compared with each other and the NCBI reference genome.** All genomes exhibited an ANI > 99.95%. The DSMZ sample displayed the highest similarity to the NCBI reference, while the Ref_WT and Ref_ETH strains were most similar to each other.

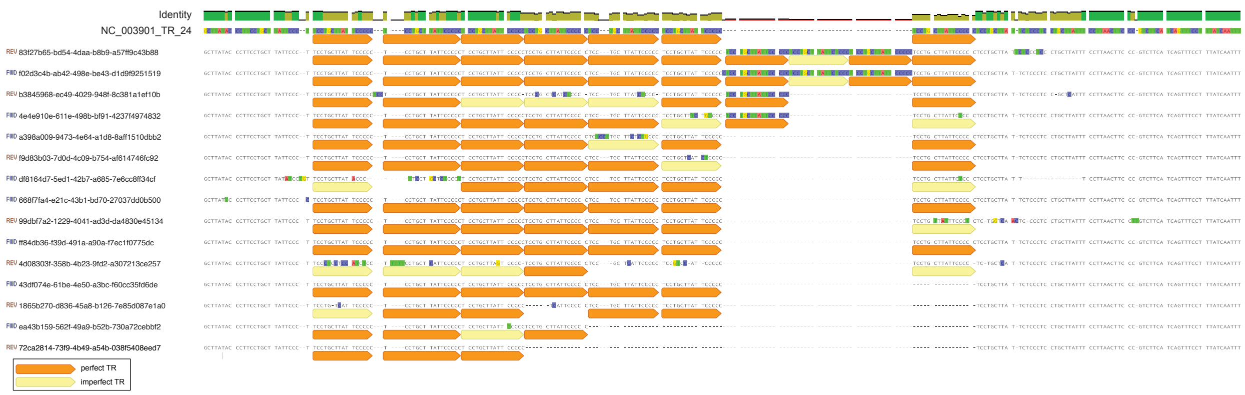

**Supplementary Fig. 5 | Oxford Nanopore long-read mapping across TR_24 analyzed using tandem-genotype.** Read-level variations span from -4 to +3 TR units relative to the reference sequence. Orange and yellow arrows indicate perfect and imperfect TR units, respectively, with the latter likely attributed to sequencing artifacts.

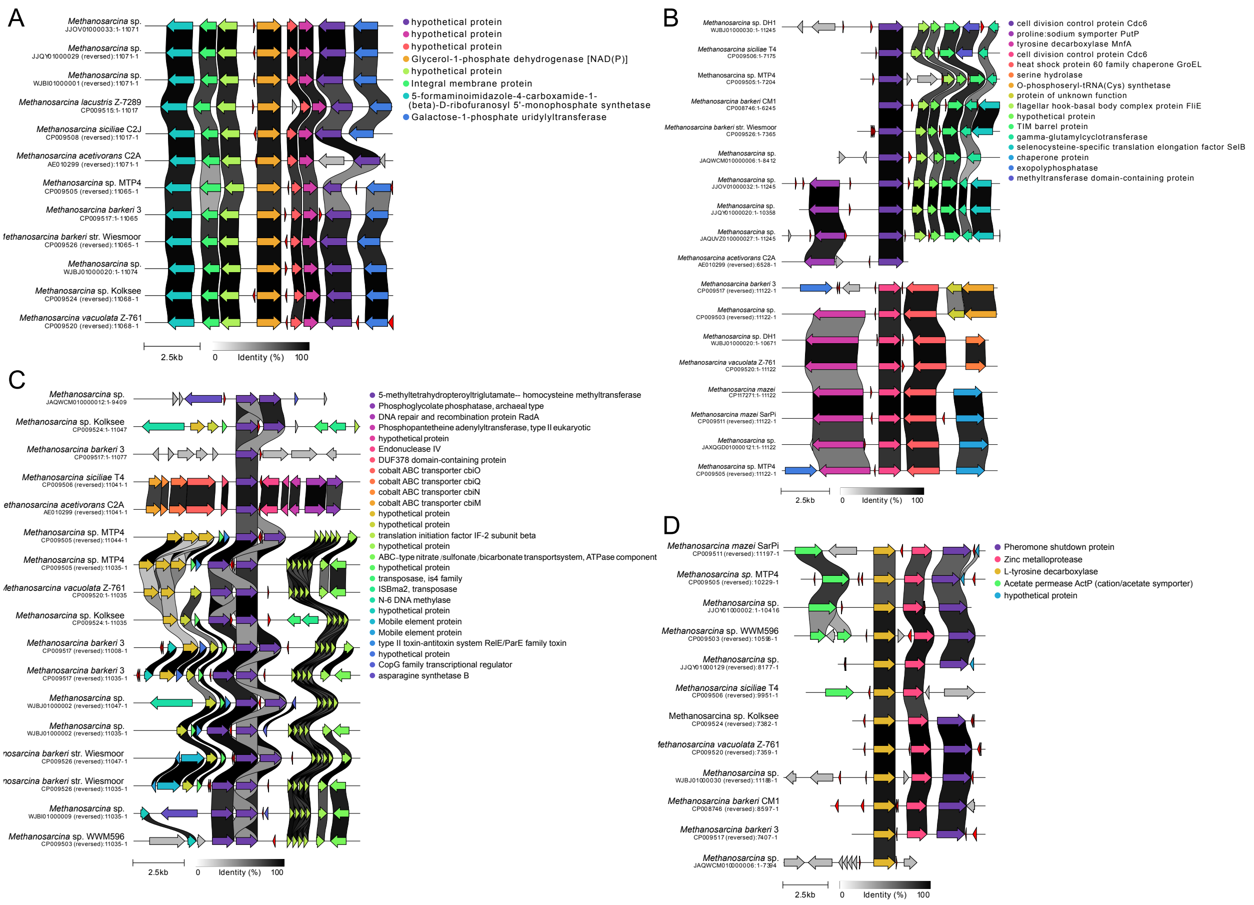

**Supplementary Fig. 6 | Genomic neighborhoods of gene clusters across *Methanosarcina* genomes where a TR (red) is frequently adjacent to a specific gene**, including the: **(A)** glycerol-1-phosphate dehydrogenase gene, **(B)** archaeal cell division control protein 6 gene, **(C)** 5-methyltetrahydropteroyltriglutamate--homocysteine methyltransferase gene, and **(D)** tyrosine decarboxylase / aspartate 1-decarboxylase gene.

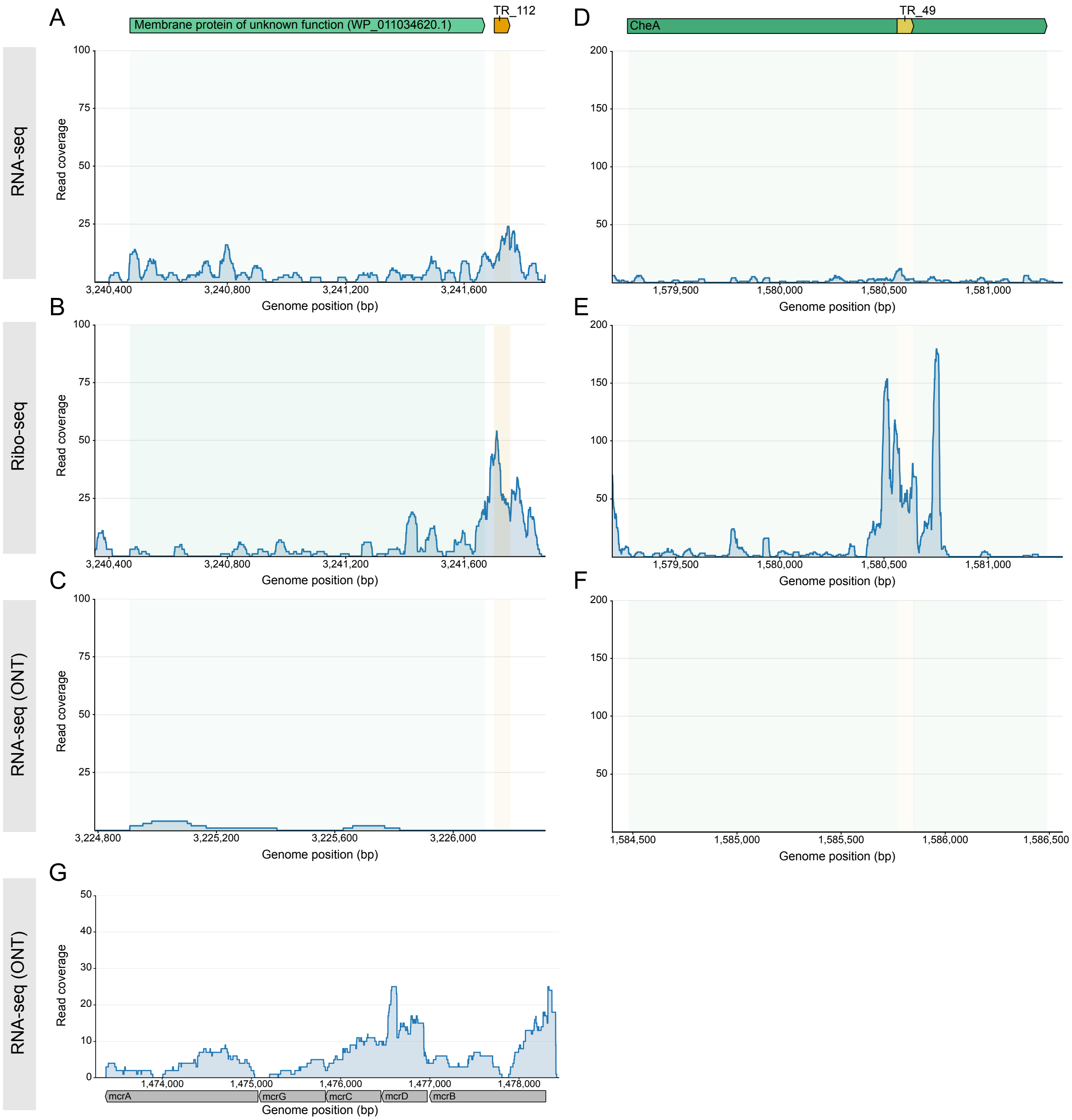

**Supplementary Fig. 7 | RNA-seq and Ribo-seq profiling of representative TR-containing loci.** Read coverage across **(A–C)** TR_112 and adjacent gene WP_011034620.1, and **(D–F)** TR_49 located within cheA (WP_011033271.1), using RNA-seq **(A,D)**, Ribo-Seq **(B,E)**, and Oxford Nanopore Technologies (ONT) RNA-Seq **(C,F)**. **(G)** ONT RNA-Seq long-read coverage across the mcr operon.

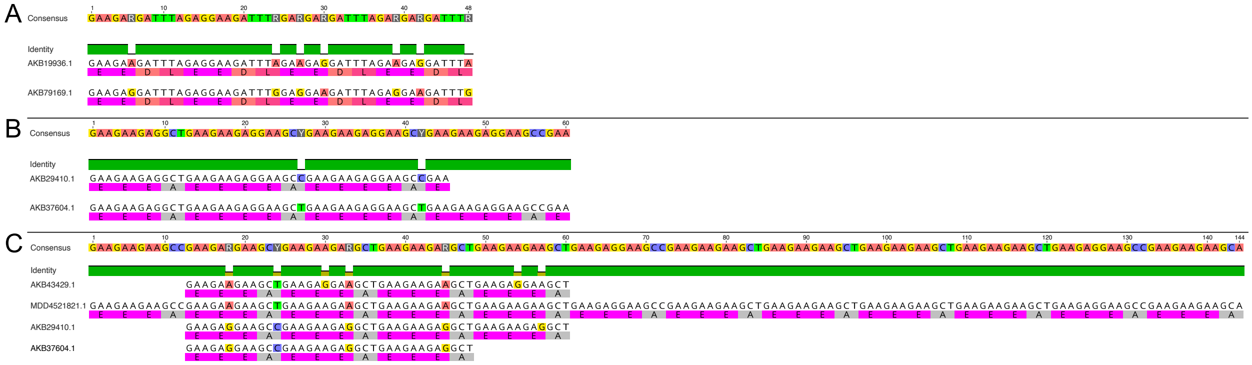

**Supplementary Fig. 8 | Sequence alignments of identical aaTRs of exosome complex RRP41 homologs in *Methanosarcina* showing variation in nucleotide sequences while conserving the amino acid repeat.** Examples are shown of the repeats [EEDL] in **(A)**, [EEEAE] in **(B)**, and [EEEA] in **(C)**.

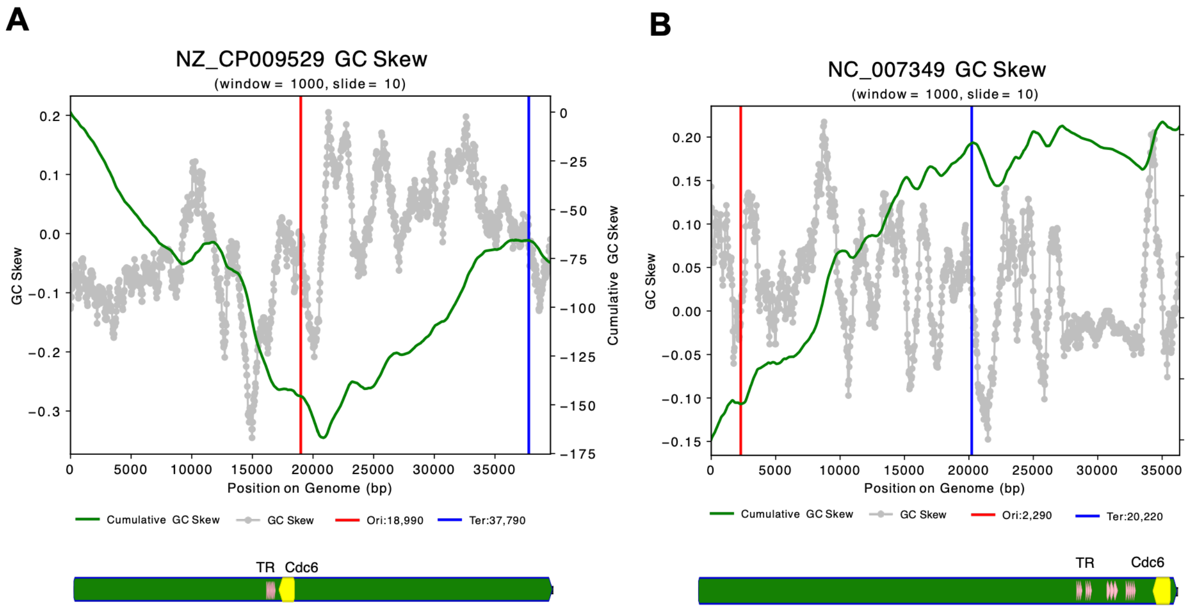

**Supplementary Fig. 9 | GC skew profiles for *Methanosarcina barkeri* plasmids (A) NZ_CP009529 and (B) NC_007349.** Below each GC skew plot, the locations of identified TR regions and the cdc6 gene are annotated. Vertical red and blue lines denote the predicted origin of replication (ori) and terminus (ter), respectively.

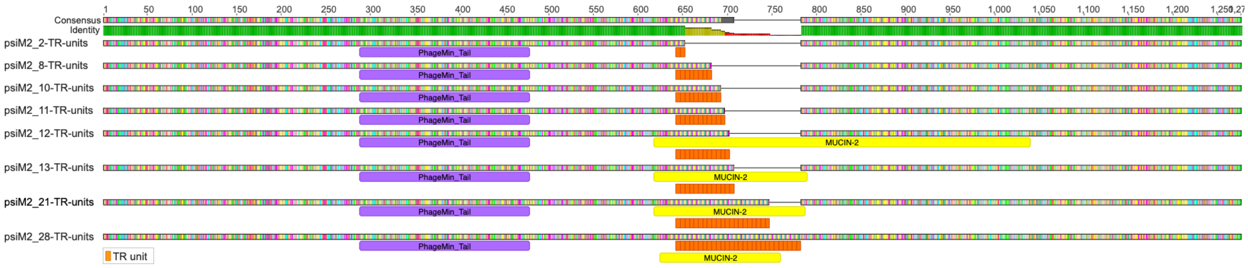

**Supplementary Fig. 10 | Different TR variants in the TMP of ΨM2 lead to the formation of a new predicted mucin-2 domain.** TR units are shown in orange.

### Supplementary Tables:

**Supplementary Table 1 | Tandem repeat densities across 50,567 prokaryotic chromosomes and complete genomes retrieved from the RefSeq database, alongside 17 curated or complete Borg genomes**. Geographic origin and ecosystem metadata are shown for the 30 *Methanosarcina* genomes.

**Supplementary Table 2.1 | Tandem repeat densities and genome statistics of 87 dereplicated (95% ANI) methanogen genomes.** Geographic origin and ecosystem metadata are also listed together with identified carbohydrate-active enzymes.

**Supplementary Table 2.2 | Methanogen genome dereplication and selection.** Summary of 252 non-dereplicated methanogen genomes and dereplication statistics at 95% ANI using dRep, alongside data availability notes and representative genome selection criteria.

**Supplementary Table 3 | *Methanosarcina* genome summary and dereplication.** Summary of 135 non-dereplicated *Methanosarcina* genomes, including genome completeness, contamination estimates, and dereplication information at a 99% ANI threshold using dRep. Tandem repeat statistics is shown for the 37 dereplicated *Methanosarcina* genomes.

**Supplementary Table 4.1 | Pairwise comparisons of dereplicated *Methanosarcina* genomes.** ANI and the proportion of shared tandem repeats across all pairwise genome combinations.

**Supplementary Table 4.2 | Identical tandem repeats across *Methanosarcina* genomes.** Includes repeat unit sequences, copy numbers, and genomic positions for all shared TRs.

**Supplementary Table 5.1 | Identical tandem repeats across long-read *M. mazei* Gö1 assemblies and the reference genome (NC_003901.1).** Includes repeat unit sequences, copy numbers, and genomic positions for all shared TRs.

**Supplementary Table 5.2 | Unique tandem repeats in long-read *M. mazei* Gö1 assemblies relative to the reference genome (NC_003901.1).** Genomic position and unit sequence are listed for each unique TR.

**Supplementary Table 6 | Single-nucleotide variants (SNVs) in long-read *M. mazei* Gö1 assemblies. Table contains two sections: (A)** Variant and SNV counts for each sample together with SNV frequency statistics. **(B)** Individual variants identified by Clair3 across all assemblies alongside corresponding quality statistics.

**Supplementary Table 7 | Enriched and depleted KEGG Orthology (KO) categories in the immediate genomic neighborhoods of intergenic tandem repeats (TRs) across *Methanosarcina* species.** The table is divided into three sections: **(A)** Significantly enriched and depleted Level C KO categories ($p_{adj}$ < 0.05) among immediate flanking genes of intergenic TRs, with empirical $p$-values derived from a 1,000-iteration non-parametric permutation test with pseudo-count and Benjamini–Hochberg FDR correction (excluded categories with < 10 observed hits). **(B–C)** Level C and higher-tier Level B KEGG categories absent from intergenic TR neighborhoods.

**Supplementary Table 8 | RNA-seq and Ribo-seq coverage across 168 tandem repeat regions of *M. mazei* Gö1.** Expression levels and local coverage profiles were categorized as follows: Low expression (approx. 5–30 reads), Moderate expression (approx. 30–100 reads), Strong expression (>100 reads), Hill (localized coverage higher than adjacent flanking regions), and Valley (localized coverage lower than adjacent flanking regions).

**Supplementary Table 9 | Top 10 predicted $sRNA_{075}$ and $sRNA_{129}$ binding partners in *M. mazei* Gö1 identified by IntaRNA.**

**Supplementary Table 10 | Enriched and depleted KEGG Orthology (KO) categories in *Methanosarcina* tandem repeat proteins (TR-proteins).** The table is divided into three sections: **(A)** Significantly enriched and depleted Level C KO categories ($p_{adj}$ < 0.05) in TR-proteins, with empirical $p$-values derived from a 5,000-iteration non-parametric permutation test with Benjamini–Hochberg FDR correction (filtering excluded categories with < 10 hits or ≤ 1 KO term); **(B)** Level C KEGG categories containing no TR-proteins together with their genomic prevalence (filtered for > 3 KO terms); and **(C)** Higher-level Level B KEGG categories containing no TR-proteins together with their genomic prevalence.

**Supplementary Table 11 | Significantly enriched protein domains in tandem repeat (TR) proteins and their structural overlap.** Statistical enrichment of functional domains and IDRs (occurring in ≥10 TR-proteins) relative to background proteins, evaluated using two-sided Fisher's exact tests with Benjamini–Hochberg FDR correction. Only significantly enriched domains (FDR < 0.05) are reported.

**Supplementary Table 12 | Cell-surface enrichment and amino acid tandem repeat co-occurrence of protein domains.** Statistical assessment of InterPro enrichment at predicted cell-surface locations (PSORTb categories: 'extracellular' and 'cell wall') across the proteomic background using two-sided Fisher's exact tests with Benjamini–Hochberg FDR correction.

**Supplementary Table 13 | Disaggregatase-related proteins across dereplicated *Methanosarcina* genomes.** Overview of 257 disaggregatase-related proteins, including predicted subcellular localizations (PSORTb) and annotated InterPro disaggregatase_repeat domains. Perfect and imperfect amino acid tandem repeats (aaTRs) were identified using a custom Python script, while nucleotide TRs were annotated using Geneious Prime (see methods section).

**Supplementary Table 14 | Tandem repeat-containing proteins (TR-proteins) in *M. ruminantium* (NC_013790.1) with perfect and degenerate amino acid TRs.** Functional annotations, predicted subcellular localizations (PSORTb), and the presence and sequences of corresponding nucleotide TRs are shown for all 43 TR-proteins.

**Supplementary Table 15 | Tandem repeat-containing proteins (TR-proteins) detected in publicly available proteomic datasets of methanogens.** Table is divided into three sections: (A) *M. mazei* Gö1 (PXD004325), (B) *M. acetivorans C2A* (PXD022016), and (C) *M. olleyae* YLM1 (PXD070174). Listed are TR-proteins (A, 10 of 22; B, 7 of 44; C, 3 of 6) detected at a protein-level FDR of 1% (protein q < 0.01) after reanalysis with Proteome Discoverer 3.3.1 and CHIMERYS (Supplementary Note 1). Columns are taken from the Proteome Discoverer protein table (the Sum PEP Score is omitted because it is infinite for most proteins); abundance and 'Found in Sample' columns are given per raw file (Fxx; file names are listed in the README of the Zenodo record). Abundances in A and B are normalized and scaled; those in C are not. In B, file F129 (181207_M42_rep3) could not be searched (Supplementary Note 1); its values were transferred from the other files by feature mapping ('Peak Found'). The only peptide lying within an aaTR that passed the 1% FDR threshold, ILELETLLVDEQR (AAM05742.1, section B), is described in the main text.

**Supplementary Table 16 | Overview of nine published methanogen plasmids.** Includes host taxonomy, plasmid sequence lengths and GC content, tandem repeat counts and statistics.

**Supplementary Table 17 | Tandem repeat region identified in the tape measure protein (AAC27060) of the ΨM2 phage (AF065411).**

**Supplementary Table 18 | Viral metagenome-assembled genomes (vMAGs) and their tandem repeat (TR) profiles.** Overview of 139 vMAGs with taxonomic classifications, predicted host taxa, genomic contexts of identified TRs, and functional predictions for TR-proteins.

**Supplementary Table 19 | Methanogen genomes excluded from this study. Overview of additional methanogen genomes published during the course of this study** and omitted from the analytical pipeline. All listed genomes harbor five or fewer tandem repeat regions.

**Supplementary Table 20 | Sources and databases for methanogen viral genomes.** List of public databases and literature-derived resources together with their source environments and number of viruses/vOTUs.

### Supplementary Note 1: Extended Methods

This Note provides detailed methods of the main text.

#### 1. Medium composition and cultivation (Organism and cultivation)

Cultures of *M. mazei* were cultivated at 37 °C without rotation in 12-ml Hungate tubes or 120-ml serum flasks containing complex medium with an $N_2$+$CO_2$ (80:20 [v/v]) gas atmosphere. The medium was prepared using anaerobic techniques[51,100] and contained: 2.0 mM $K_2HPO_4 \times 3H_2O$, 1.67 mM $KH_2PO_4$, 1.7 mM $CaCl_2 \times 2H_2O$, 2.03 mM $MgSO_4 \times 7H_2O$, 7.2 µM $FeSO_4 \times 7H_2O$, 38.5 mM NaCl, 11.0 mM Na-acetate × $3H_2O$, 9.35 mM $NH_4Cl$, 2 g/L yeast extract, 2 g/l casitone, 0.3 ml/l trace element solutions SL 6, 1 ml/l Wolin's vitamin solution, 1 ml/l resazurin and were buffered with 10.1 mM $NaHCO_3$ (modified DSM medium 120c). The pH was adjusted to 6.9. Prior to inoculation, the medium was supplemented with L-cysteine and $Na_2S$ as reducing agents (0.37 mM and 0.38 mM, respectively), and with methanol and trimethylamine as energy and carbon sources (75 mM and 20 mM, respectively). Growth was determined by measuring the optical density at 600 nm ($OD_{600}$).

#### 2. Mismatch-tolerant tandem repeat detection

Perfect tandem repeats were identified with a custom Python script, which builds on MUMmer v3.23[60] as previously described (https://github.com/rohansachdeva/tandem_repeats)[25], using the thresholds given in the main text (nucleotide TRs: no mismatch, total region length ≥ 50 nt, ≥ 3 repeat units; amino acid TRs: ≥ 16 aa region length, ≥ 3 TR units; degenerate protein repeats: ≥ 9 aa region length and ≥ 3 TR units). To evaluate how many imperfect TRs are excluded from this analysis, the script was manually modified to allow mismatches (up to two per repeat unit). The initial exact seed size used to nucleate candidate arrays is reduced dynamically to account for point mutations, and candidate arrays are then extended by comparing each adjacent repeat unit against the seed unit using Hamming distance, accepting units with varying number of mismatches. We allowed one mismatch for *M. mazei*, as even a single mismatch flagged a substantially larger set of degenerate TRs, and two mismatches to detect additional TR-containing cell-surface proteins in *M. ruminantium*. All other parameters, filters and output formats are identical to the perfect-repeat search, so that perfect and mismatch-tolerant calls can be compared directly for the same genome.

#### 3. Negative binomial mixed model of TR density and sensitivity analysis

To identify drivers of TR density in 170 methanogen genomes, a multivariable Negative Binomial Generalized Linear Mixed-Effects Model was fitted to raw TR counts with glmmTMB in R[101,102]. Fixed effects were log-transformed genome size, broad ecosystem classification, the optical growth conditions predicted with GenomeSPOT and the methanogenesis pathways encoded according to DRAM. Genus identity modeled as a random intercept. The significance of fixed effects significance was evaluated using Type II Wald Chi-squared ($\chi^2$) tests, and the random intercept was evaluated via Likelihood Ratio Test against the model without the random term. To evaluate parameter stability and to rule out bias from unequal sample sizes across habitats (n = 8–34 genomes per group), two stratified bootstrapping schemes were run for 100 iterations each (native group ratios vs. 8 genomes per group). Estimates and significance of the fixed effects were compared between the full model and the bootstrap replicates. The enrichment of TRs in *Methanosarcina* genomes relative to all other methanogen genera was further tested using a Wilcoxon rank-sum test.

##### 4. Functional enrichment in TR neighborhoods

Genomes of *Methanosarcina* spp. were dereplicated to 99% ANI, resulting in 28 genomes with predicted coding sequences; only intergenic TRs were included in this analysis. For each genome, ORFs were characterized using KofamScan v1.3.0[62] against the KEGG Orthology (KO) database (ko00001; accessed March 10, 2026) and filtering for the top-scoring hit per ORF (*E*-value < 0.001). The direct genomic neighborhood of each TR was defined as the immediate upstream and downstream flanking genes. To determine if specific functional groups and their higher hierarchical categories (KO level C) were significantly enriched or depleted within these neighborhoods, an empirical, non-parametric permutation test was performed over 1,000 iterations. The observed gene counts within the TR neighborhoods were compared against a null distribution generated by randomly sampling an equivalent number of genes from the annotated background. To ensure robust statistical power, functional categories with fewer than 10 observed hits and categories with only one KO were excluded from the analysis. Two-sided empirical *p*-values were calculated incorporating a pseudo-count correction[103,] deviations from the background were quantified using permutation-derived *Z*-scores[104], and multi-testing corrections were applied using the Benjamini-Hochberg FDR correction[63].

Functional categories that were present in the background proteome but completely absent from TR neighborhoods were identified and characterized by their absolute protein counts, relative abundance within each annotation level, and associated KO composition. However, because these functional categories consisted mostly of only one to three genes, they were not further investigated.

To determine if specific functional genes are significantly enriched or depleted within the genomic TR neighborhood, an exact two-sided Binomial test was applied to each individual KEGG Ortholog category[103]. For each KO, the test evaluated the number of functional gene copies located in the direct TR-neighborhood relative to the total number of copies found across the entire background dataset. The expected baseline probability was defined as the "global TR density", representing the total proportion of all annotated genes located next to TRs in the background dataset (7.26%). KOs with neighborhood proportions significantly higher than the global TR density were classified as enriched.

##### 5. Statistical characterization of aaTR proteins

Subcellular localization was predicted with PSORTb v3.0[64] in archaeal mode; proteins with an 'unknown' localization assignment (14.0% of the global proteomic background and 28.8% of TR-proteins) were excluded from downstream analyses. Enriched localizations within the TR-proteins were identified relative to the global proteomic background using a two-sided Fisher's Exact Test. To accommodate low-abundance localizations, *p*-values were computed via Monte Carlo simulation (10,000 replicates), and effect sizes were evaluated by calculating post-hoc sample odds ratios for each localization against all alternative categories combined[104]. Amino acid sequences of TR-proteins were profiled using KofamScan v1.3.0[62] and intrinsic disordered regions (IDRs) were predicted with MobiDBLite v2.0[65]. These functional annotations were further validated or curated using InterProScan v5.69-101.0[66].

Differences in TR-protein length and TR coverage percentage across subcellular localization groups were evaluated using Kruskal-Wallis[107] tests with post-hoc Dunn's tests[108], while multi-TR protein prevalence was assessed using Chi-Square and pairwise Fisher's exact tests[109]. Variations in TR amino acid composition across sub-cellular localizations were determined using a Chi-squared ($\chi^2$) test of independence[110], with specific enrichment and depletion patterns identified from standardized residuals following Benjamini-Hochberg FDR correction[63]. To evaluate TR composition biases, the relative abundance of each amino acid within aaTR regions was compared against the entire proteomic background of each proteome using a relative difference metric, with significance determined via a two-sided, paired Wilcoxon signed-rank test[109] with Benjamini-Hochberg FDR correction[63]. Concurrently, a background presence/absence matrix was constructed for IDRs and the top 25 most frequent InterPro domains. Enrichment of these domains in TR-protein was assessed using two-sided Fisher's Exact Tests with subsequent FDR correction.

For each unique protein the amino acid coordinates of functional domains and IDRs were mapped against TR boundaries to calculate how much they overlapped. Protein domains were then grouped into three categories: TR-driven (where the TR makes up at least 50% of the domain), partial overlap (less than 50% overlap), or separate elements that are just encoded on the same protein. To identify domains that are preferentially located at the cell surface and that co-occur with aaTRs, unique InterPro domains were evaluated for spatial enrichment across the proteomic background. Two-sided Fisher's Exact Tests[107] were applied to each domain to assess non-random association with predicted cell-surface localization (PSORTb categories 'extracellular' and 'cell wall'), quantifying effect sizes via sample odds ratios with *p*-values adjusted for multiple testing[63]. Domains passing the surface-enrichment test were then intersected with the set of aaTR-containing proteins to determine co-occurrence frequency (Supplementary Table 12).

To determine whether specific functional proteins and their higher hierarchical categories (KO level C) were significantly enriched or depleted within TR-proteins, an empirical, non-parametric permutation test was performed over 5,000 iterations across the annotated KEGG Orthology hierarchy. The observed TR-protein counts were compared against a null distribution generated by randomly sampling an equivalent number of proteins from the annotated background proteome to compute permutation-derived *Z*-scores[104] and two-sided empirical *p*-values with Benjamini-Hochberg FDR correction[63]. To ensure robust statistical power, functional categories with fewer than 10 observed hits and categories with only one KO were excluded from the analysis. Higher functional categories that were completely absent from the TR-protein pool were isolated and profiled by their absolute copy-number and number of individual KOs (filtered for $n > 3$) within the global background proteome.

#### 6. Nanopore whole-genome sequencing

Cells of *M. mazei* DSMZ and Ref_ETH cultures were harvested during exponential growth by centrifugation at 8,000 g for 5 min with downstream genomic DNA isolation using Qiagen's MagAttract HMW DNA Kit and *M. mazei* Ref_WT by centrifugation at 13,000 g for 2 min and genomic DNA isolation using the Wizard Genomic DNA Purification Kit (Promega) following the manufacturers' protocols. Genomic DNA was purified using AMPure XP beads and libraries were prepared using the Native Barcoding Kit V14 following the protocol for genomic samples (SQK-NBD114.24; version NBE_9169_v114_revU_30Jan2025). DNA was sequenced using the Oxford Nanopore PromethION 2 Solo system to a sequencing depth of 275,000–2,190,000 reads. Data produced was basecalled using the model dna_r10.4.1_e8.2_400bps_sup@v5.0.0 with Dorado v7.8.3. Raw reads were filtered for Q-score ≥ 15 and ≥ 1,000 bp using chopper v0.10.0[68]. An additional filtering using Filtlong (https://github.com/rrwick/filtlong) was performed with adjusted parameters (--target_bases 1Gb, 400Mb or 210Mb) depending on the read quality and sequencing depth of each sample. Genomes were assembled using the reference-based assembler Rebaler v0.2.0 (https://github.com/rrwick/rebaler) using NC_003901.1 as reference. The TR regions were identified in the assembled genomes as described above and ORFs were transferred from the reference genome to the assemblies using liftoff v1.6.3[112].

#### 7. Short-read RNA-seq and Ribo-seq processing

Ribosome profiling (Ribo-seq) and RNA sequencing (RNA-seq) datasets were downloaded from the NCBI Sequence Read Archive (SRA) on 13 July 2026 under BioProject accession PRJNA1004312 (GEO series GSE240615)[7]. Adapter sequences were trimmed and low-quality reads were removed using Trimmomatic v0.39[74] in single-end mode ('ILLUMINACLIP:TruSeq3-SE.fa:2:30:10 LEADING:3 TRAILING:3 SLIDINGWINDOW:4:15'). Ribosomal RNA reads were subsequently filtered using Bowtie2 v2.5.4[75] ('--very-sensitive-local') mapped against the SortMeRNA database v4.3[76]. High-quality clean reads were then aligned to the reference genome (NC_003901.1) using Bowtie2 with the same parameters, and genome coverage was analyzed using Geneious Prime 2026.1.1 (https://www.geneious.com). For each TR region the expression was categorized into "low" (5–29 reads), "moderate" (30–100 reads), and "high" (> 100 reads).

#### 8. Long-read transcriptome sequencing

For validation of TR expression using long-read sequencing, *M. mazei* Gö1 cultures were harvested during exponential growth ($OD_{600}$ ~ 0.2) by centrifugation at 13,000 g for 2 min. Total RNA was extracted using RotiZol and DNaseI treated as described previously[77]. Ribosomal RNA was depleted following siTOOLs Pan-Archaea riboPOOL kit (dp-K012-27) with subsequent ethanol precipitation. rRNA depleted RNA samples were polyA tailed following the Oxford Nanopore protocol "3' polyadenylation of RNA transcripts using *E. coli* poly(A) polymerase (PAP)" v5. Libraries were prepared using the cDNA-PCR Barcoding Kit V14 (SQK-PCB114.24) and were sequenced using the Oxford Nanopore PromethION 2 Solo system to sequencing depths of 175,327 (41,527,892 bases) and 567,357 reads (102,537,092 bases). After base calling using the model dna_r10.4.1_e8.2_400bps_sup@v5.0.0 with Dorado v7.9.8, raw reads were length filtered for ≥ 15 bp using chopper v0.10.0[68], and artificially added 3' polyA tails were removed using cutadapt v4.4[78] with following parameter: "-a A{10} -e 3 -j 0". Reads were quality filtered for Q-score ≥ 10 before remaining rRNA reads were removed using minimap2 v2.22[72] ("-ax splice -uf") with the sensitive SortMeRNA database[76] together with the known rRNA sequences of the *M. mazei* Gö1 (NC_003901.1) genome. Non ribosomal reads were extracted from the alignment using samtools v1.17[79], and mapped to the assembled genome using minimap2 v2.22 with the same parameters as listed above. Mapped reads were analyzed using Geneious Prime 2026.1.1 (https://www.geneious.com).

#### 9. Proteomics search parameters

All vendor-format raw mass spectrometry files from *M. mazei* Gö1 (PXD004325; 72 files), *M. acetivorans* C2A (PXD022016; 15 files), and *Methanobrevibacter olleyae* YLM1 (PXD070174; 10 files) were retrieved from the PRIDE database[80] and reanalyzed with the same search settings for all three datasets in Proteome Discoverer 3.3 (v3.3.1.110) using CHIMERYS[81] (v5.2.10) with the INFERYS 5.8 fragmentation model. One PXD022016 file (181207_M42_rep3) could not be searched by CHIMERYS because one of its spectra (scan 43417) contains no peaks; PXD022016 identifications are therefore based on 14 of the 15 files. Searches used fully specific Trypsin/P digestion with up to two missed cleavages, peptide lengths of 7–30 amino acids, precursor charge states of 1–6, static cysteine carbamidomethylation, dynamic methionine oxidation (at most three dynamic modifications per peptide), and a maximum fragment-ion mass tolerance of 20 ppm. Peptide-spectrum matches (PSMs) and peptides with global q-values below 0.01, corresponding to a 1% FDR threshold, were classified as high confidence and retained for downstream analysis. Proteins supported by at least one retained peptide were grouped automatically using strict parsimony and evaluated at a protein-level FDR threshold of 1%. Only proteins with q-values below 0.01 were counted as detected proteins and used to calculate detection rates. Searches used organism-specific whole-proteome protein databases, supplemented with labeled duplicate entries for the TR-proteins (22, 44 and 6, respectively; identical in sequence to whole-proteome entries) and, for *M. mazei* Gö1, 69 putative small TR-protein sequences, together with a common-contaminant protein-sequence database of 244 unique protein sequences[82].

##### 10. Viral genome dataset and annotation

Viral genome sequences were compiled from multiple public databases and literature-derived resources (Supplementary Table 20) targeting methanogen hosts, generating an initial dataset of 447,591 viral genomes spanning diverse environments. Sequences were dereplicated using seqkit v2.12.0[85] using rmdup -s. The resulting 407,171 non-redundant viral genomes were screened for tandem repeats as described above, identifying 18,510 TR loci across 15,889 viral genomes. GeNomad v1.8.1[86] was used to assign viral taxonomy and confirm that all sequences were classified as viral. Protein-coding genes were predicted using Prodigal-gv v2.11.0[87], and genome annotation was performed using Pharokka v1.9.1[88], eggNOG-mapper v2.1.12[89], KofamScan v1.3.0[62], InterProScan v5.69-101.0[66], and Colabfold v1.5.5[91] together with FoldSeek v8-ef4e960[91] (against PDB released January 1, 2024) to improve functional characterization of viral proteins using a cutoff e-value of $10^{-4}$. Annotations were manually assessed and grouped into functional categories (Supplementary Table 18). Putative host associations were inferred using iPHoP v1.4.2[92], from which viruses predicted to infect methanogenic archaea were selected for downstream analysis. Genome quality was evaluated using CheckV v1.0.3[93], retaining genomes with ≥ 50% completeness and at least two predicted genes. TR loci were categorized according to their genomic position relative to predicted ORFs.
